# Single-cell epigenomic dysregulation of Systemic Sclerosis fibroblasts via CREB1/EGR1 axis in self-assembled human skin equivalents

**DOI:** 10.1101/2024.03.22.586316

**Authors:** Tamar R. Abel, Noelle N. Kosarek, Rezvan Parvizi, Helen Jarnagin, Gretel M. Torres, Rajan Bhandari, Mengqi Huang, Diana M. Toledo, Avi Smith, Dillon Popovich, Michael P. Mariani, Heetaek Yang, Tammara Wood, Jonathan Garlick, Patricia A. Pioli, Michael L. Whitfield

**Affiliations:** Department of Biomedical Data Science, Geisel School of Medicine at Dartmouth, Lebanon, NH, USA; Department of Molecular and Systems Biology, Geisel School of Medicine at Dartmouth, Hanover, NH, USA; Department of Diagnostic Sciences, Tufts University School of Dental Medicine, Boston, MA, USA; Department of Microbiology and Immunology, Geisel School of Medicine at Dartmouth, Lebanon, NH, USA; Division of Rheumatology and Clinical Immunology, Department of Medicine, School of Medicine, University of Pittsburgh, Pittsburgh, PA, USA; Department of Epidemiology, Geisel School of Medicine at Dartmouth, Lebanon, NH, USA

## Abstract

Systemic sclerosis (SSc) is an autoimmune disease characterized by skin fibrosis, internal organ involvement and vascular dropout. We previously developed and phenotypically characterized an *in vitro* 3D skin-like tissue model of SSc, and now analyze the transcriptomic (scRNA-seq) and epigenetic (scATAC-seq) characteristics of this model at single-cell resolution. SSc 3D skin-like tissues were fabricated using autologous fibroblasts, macrophages, and plasma from SSc patients or healthy control (HC) donors. SSc tissues displayed increased dermal thickness and contractility, as well as increased α-SMA staining. Single-cell transcriptomic and epigenomic analyses identified keratinocytes, macrophages, and five populations of fibroblasts (labeled FB1 – 5). Notably, FB1 APOE-expressing fibroblasts were 12-fold enriched in SSc tissues and were characterized by high EGR1 motif accessibility. Pseudotime analysis suggests that FB1 fibroblasts differentiate from a TGF-β1-responsive fibroblast population and ligand-receptor analysis indicates that the FB1 fibroblasts are active in macrophage crosstalk via soluble ligands including FGF2 and APP. These findings provide characterization of the 3D skin-like model at single cell resolution and establish that it recapitulates subsets of fibroblasts and macrophage phenotypes observed in skin biopsies.

## Introduction

Systemic sclerosis (SSc) is a rare systemic autoimmune disease characterized by immune dysregulation, skin fibrosis, organ dysfunction, and vascular abnormalities (1). SSc has the highest case fatality rate among all autoimmune diseases, with approximately 30% of patients dying within ten years of diagnosis (2). There are only two FDA-approved therapies, nintedanib and tocilizumab, both for the treatment of SSc interstitial lung disease (3, 4). The development of much-needed new treatments to combat SSc could be accelerated with the use of *in vitro*, pre-clinical models constructed from human cells that recapitulate key disease phenotypes and complement existing drug-testing strategies.

Single-cell transcriptomic studies in SSc skin have provided valuable insight into SSc pathogenic cell types and molecular drivers (5–7). In particular, multiple subtypes of fibroblasts have been identified, with specific populations showing increases or decreases in SSc skin. However, the application of skin biopsies to pre-clinical studies is limited in sustainability due to the difficulty of obtaining skin biopsies from patients with a rare disease.

To address this issue, we developed a validated *in vitro* 3D tissue model system that allows us to fabricate human skin equivalents from human patient-derived cells and plasma (8). This model, named self-assembled skin equivalent (saSE), recapitulates many of the genotypic and phenotypic characteristics of SSc patient skin, including increased tissue thickness and stiffness. As a result, their complex microenvironment holds many advantages over conventional monolayer culture when studying SSc patient-derived cellular subsets at the single cell level. Beyond this, our 3D tissue model significantly differs from other commonly used 3D models in that it does not use any non-human components, such as bovine collagen as extracellular matrix. Since fibroblasts and macrophages have been implicated in regulation of SSc skin fibrosis, this tissue model allows us to focus on the complex interactions between a subset of the cell populations most likely to result in identification of molecular targets of interest in scleroderma (9–11).

While several studies have implicated epigenetic changes in fibroblasts in disease pathogenesis and associated fibrosis (12–14), relatively little is known about epigenetic changes in SSc fibroblasts at single-cell resolution. Recent single-cell analyses in SSc dermal fibroblasts have focused on transcriptional profiles, with inferred epigenetic regulation or epigenetic analyses on only a small population of cells (5, 7). One study performed bulk Assay for Transposase Accessible Chromatin with sequencing (ATAC-seq) analyses on specific cell-types isolated by flow-sorting (15). While providing valuable insights, this approach may not fully capture the breadth of within-cell type heterogeneity.

To address this, we leveraged our saSE model using single-cell transcriptomic and epigenomic data to recapitulate *in vitro* cellular subsets that have been observed in SSc skin *in vivo*. We identified five fibroblast populations (labeled FB1 – 5) and mapped them to populations found in human skin. The FB1 fibroblast subpopulation was enriched in SSc saSE 3D tissues and was characterized by dysregulation of EGR1 transcription factor motif accessibility. Ligand-receptor (L-R) analyses predicted that FB1 fibroblasts participate in macrophage-fibroblast crosstalk through expression of soluble ligands FGF2 and APP, which are predicted to induce *CTHRC1* expression in macrophages. Pseudotime analyses suggest that the FB3 population, characterized by expression of TGF-β1 response pathways, may be the precursor of the FB1 population. Differentiation was predicted to occur via a transcription factor cascade including HIF1A, CREB1, and EGR1. Based on these results, we propose a model whereby TGF-β1 signaling in FB3 fibroblasts, differentiation of FB1 fibroblasts, and subsequent L-R interactions with macrophages creates a pro-fibrotic positive feedback loop, contributing to chronic fibrosis in SSc skin.

## Results

### Self-Assembled Skin Equivalent Tissues Recapitulate SSc Skin *in Vitro*

saSE tissues were constructed by seeding fibroblasts and monocytes, which differentiate into macrophages, together in a Transwell in media that stimulates ECM deposition to form a dermal layer **(Fig. 1A).** At the beginning of week two, tissues were exposed to SSc or healthy control (HC) plasma for seven days. At week three, normal human keratinocytes (NHKs) derived from newborn human foreskin were layered over the dermis to establish a well-differentiated stratified epithelium. The procedure and initial characterization of the saSE system at a gross phenotypic level has been previously described (8).

**Figure 1.**
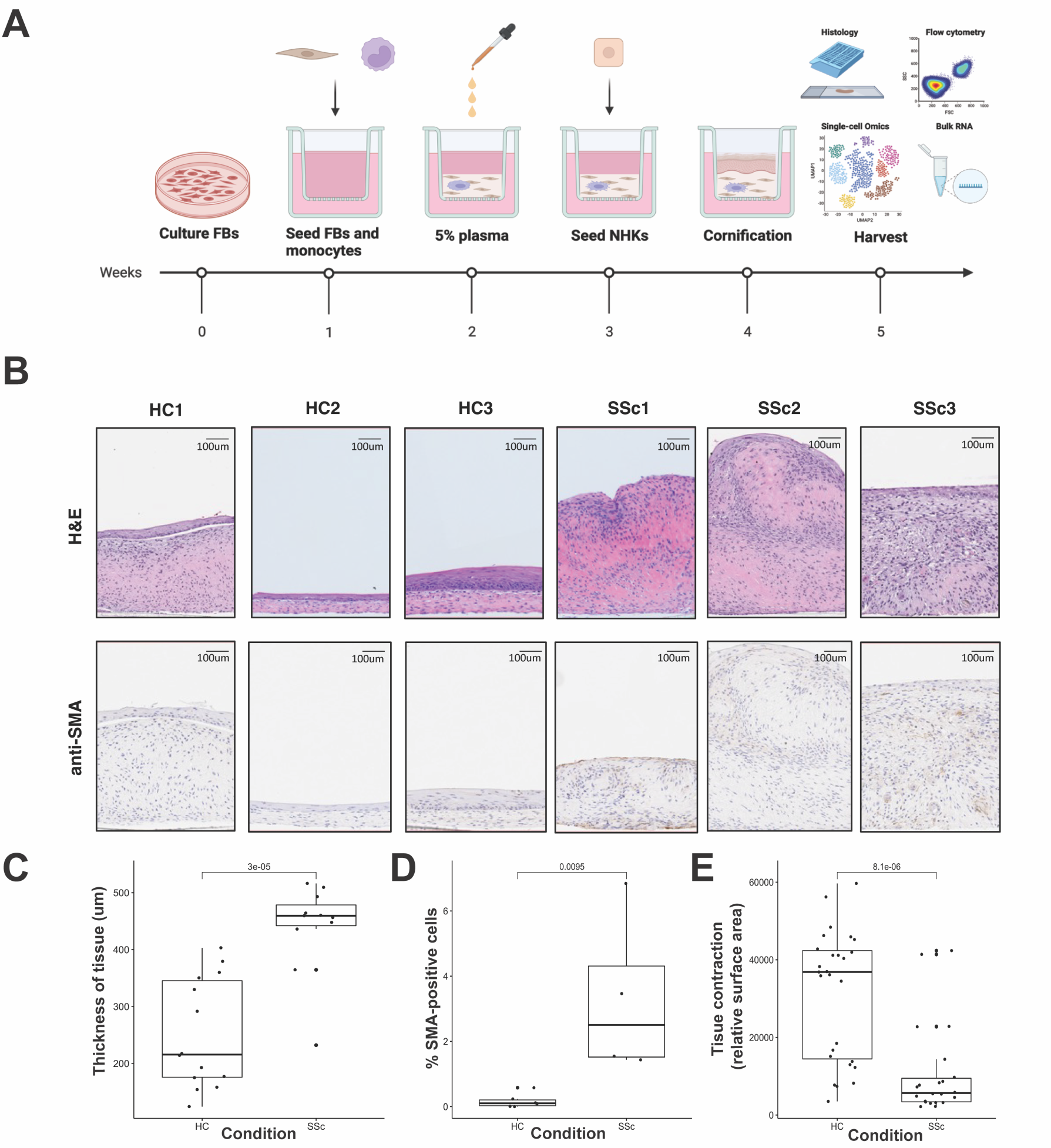
SaSE tissues recapitulate SSc fibrosis in vitro. A) An illustrated overview of Self-assembled Skin Equivalent (saSE) tissue fabrication process. B) Representative histological images of saSE tissue cross-sections stained with H&E and anti-SMA antibody for HC (n=3) and SSc (n=3) conditions. Scale bar of 100um is included in top righthand corner of each histological section. C) Average thickness of tissues based on quantification from histological images. D) Percent of SMA-positive cells based on quantification from anti-SMA stained sections. E) Surface area of tissues based on quantification from overhead images (see Figure S1 for representative images). FBs = fibroblasts, NHKs = normal human keratinocytes, HC = healthy control, SSc = systemic sclerosis, saSE = self-assembled skin equivalent

Tissues were generated using autologous fibroblasts, macrophages, and plasma from three healthy HC and three SSc donors and foreskin NHKs (cell donor information Table S1). Consistent with prior findings (8, 16), the SSc tissue histology showed a thicker dermis as visualized by H&E staining (**Fig. 1B)**. Tissue thickness was quantified by measuring three different points for each tissue in multiple histological sections and demonstrated a statistically significant increase in thickness in SSc tissues compared with HC (**Fig. 1C**; p = 3e-05). We also observed increased numbers of χξ-SMA expressing cells in SSc samples compared to HC tissues **(Fig 1D**; p = 0.0095), and tissues grown from SSc patient cells were markedly more contracted **(Fig. 1E**; Fig. S1; p = 8.1e-06).

### Single-cell RNA-seq Reveals Clusters with Disease-Specific Enrichment

scRNA-seq data were filtered to remove low quality cells, log-normalized, integrated, and scaled using regression to correct for batch effects (Fig. S2A-C). Clustering resulted in a total of seven clusters including macrophages, NHKs, and five distinct clusters of fibroblasts **(Fig. 2A**, Fig. S3A, Table S2). Major cell types were identified by enrichment of cell-specific gene markers for keratinocytes (*KRT6A, KRT17*), macrophages (*C1QB, CD163*), and fibroblasts (*VIM, THY1*) (**Fig. 2B**). Fibroblast subclusters were further defined by top differentially expressed genes.

**Figure 2.**
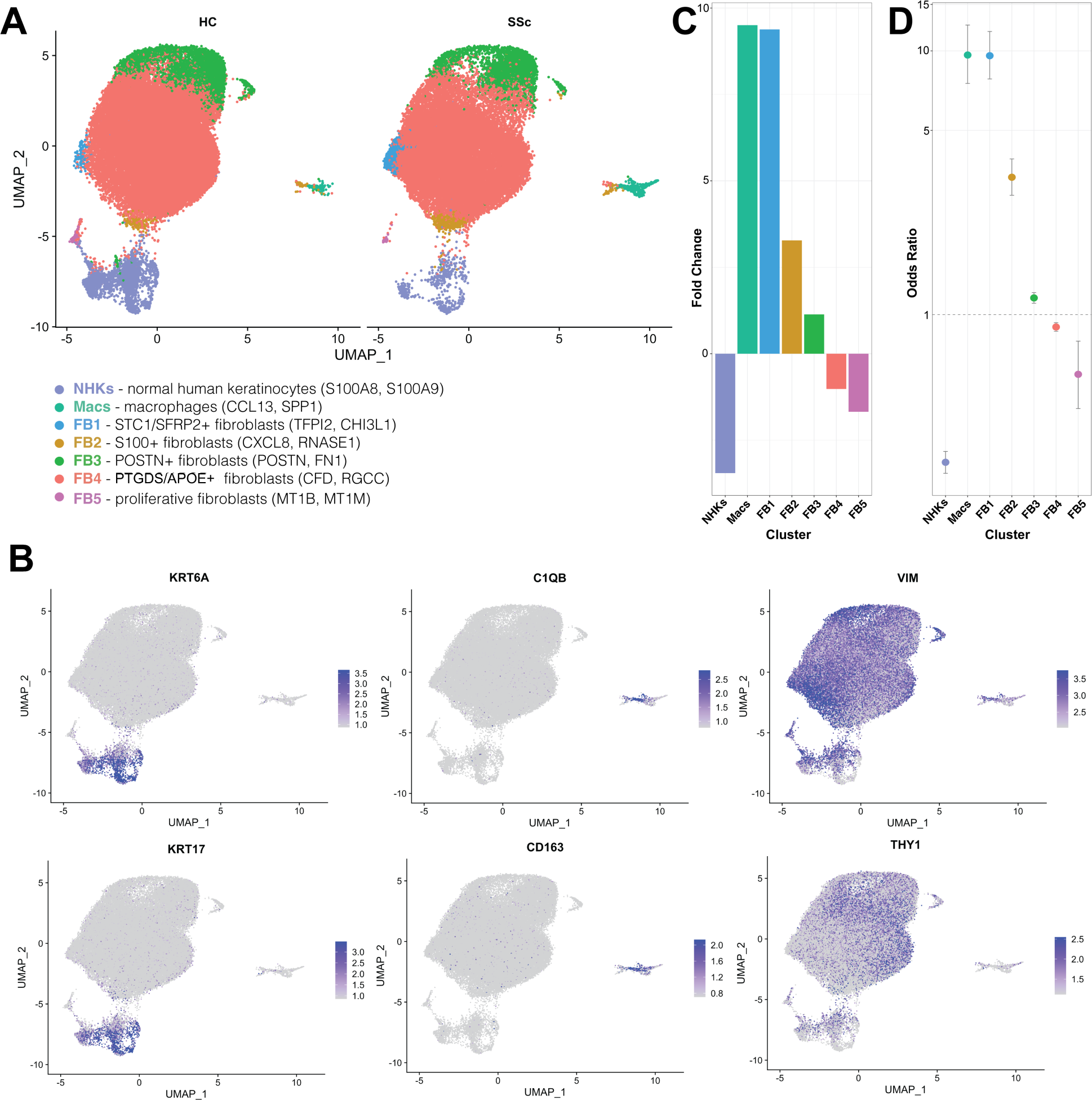
Clustering and differential enrichment of unique cell populations in single-cell RNA-seq data from saSE tissues. B) UMAP projection of integrated scRNA-seq data split by condition (HC or SSc). Clusters include normal human keratinocytes (NHKs), macrophages (Macs), and fibroblasts including five fibroblast subclusters (FB1, FB2, FB3, FB4, and FB5). Parentheses following cell cluster name include top two differentially expressed genes for that cluster. A) Feature plot of integrated scRNA-seq data with projected gene expression of cell-type specific genes for keratinocytes *(KRT6A, KRT17),* macrophages (*C1QB, CD163*), and fibroblasts (*VIM, THY1*). C) Fold-change enrichment and odds ratio (D) for SSc tissue clusters. HC = healthy control, SSc = systemic sclerosis, saSE = self-assembled skin equivalent

Fold change (FC) enrichment and odds ratio (OR) were calculated to determine proportional differences between SSc and HC tissues **(Fig. 2C-D**, Table S3-4). Macrophages (FC = 9.51), FB1 fibroblasts (FC = 9.38), and FB2 fibroblasts (FC = 3.28) were enriched in SSc samples **(Fig. 2C**, Table S3). FB3 fibroblasts were marginally enriched (FC = 1.14), while fibroblasts in clusters FB4 and FB5 were less prevalent (FC = −1.02 and −1.68) in SSc tissue. Notably, SSc tissues were significantly more likely to contain macrophages (OR = 9.65, 95% CI [7.52, 12.53]), FB1 fibroblasts (OR = 9.59, 95% CI [7.83, 11.85]) and FB2 fibroblasts (OR = 3.32, 95% CI [2.84, 3.89]) than HC tissues (**Fig. 2D**, Table S4).

### Single-cell ATAC-seq Identifies Epigenetically Distinct Fibroblasts

Similarly, scATAC-seq data were filtered to remove low quality cells, normalized, and integrated (Fig. S2D-F). Clustering analysis using the clustree package showed seven distinct clusters including macrophages, NHKs, and five clusters of fibroblasts **(Fig. 3A**, Fig. S3F, Table S1), paralleling the results obtained with scRNA-seq. Major cell types were identified using feature plots to identify populations with enriched accessibility of cell-specific transcription factor (TF) motifs for keratinocytes (SP1, TFAP2A), macrophages (SPI1, SPIC), and fibroblasts (TWIST1, NFIX) (**Fig. 3B**) (17–19). Notably, the FB1 fibroblast cluster maintained a distinct epigenetic profile from other fibroblast populations **(Fig. 3A).**

**Figure 3.**
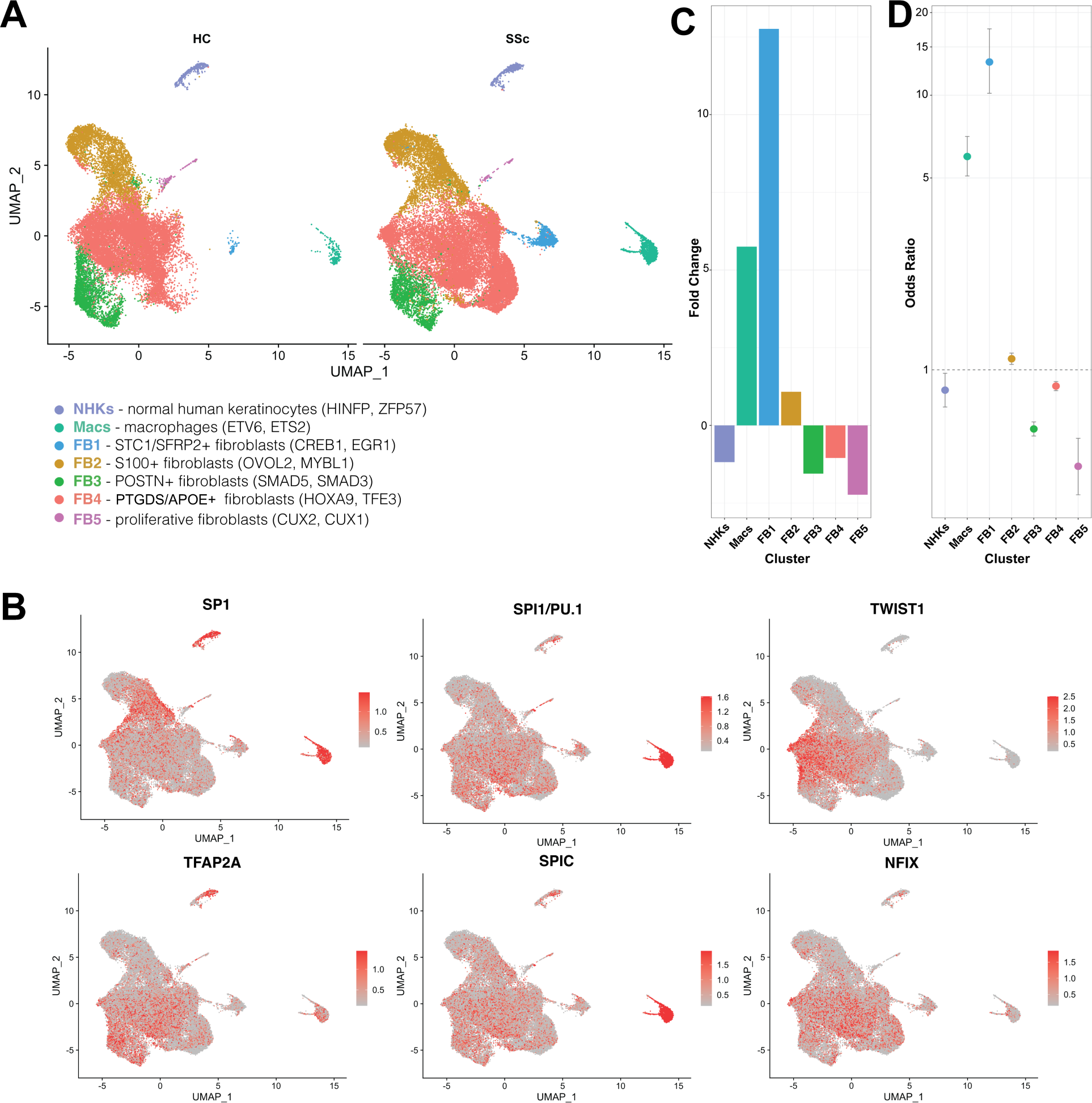
Clustering and differential enrichment of unique cell populations in single-cell ATAC-seq data from saSE tissues. B) UMAP projection of integrated scATAC-seq data split by condition (HC or SSc). Clusters include normal human keratinocytes (NHKs), macrophages (Macs), and fibroblasts including five fibroblast subclusters (FB1, FB2, FB3, FB4, and FB5). Parenthesis following cell cluster name include top two differentially accessible transcription factor motifs for that cluster. B) Feature plots of integrated scATAC-seq data with projected motif accessibility of cell-type specific transcription factors for keratinocytes (SP1, TFAP2A), macrophages (SPI1, SPIC), and fibroblasts (TWIST1, NFIX). C) Fold-change enrichment of cell cluster proportions and C) Fold-change enrichment and odds ratio (D) for SSc tissue clusters. HC = healthy control, SSc = systemic sclerosis, saSE = self-assembled skin equivalent.

Consistent with scRNA-seq results, fold enrichment analysis of scATAC-seq data showed an enrichment of macrophages (FC = 5.75), FB1 fibroblasts (FC = 12.76), and FB2 fibroblasts (FC = 1.10) **(Fig. 3C**, Table S2) in SSc tissues, while clusters FB4 and FB5 were less prevalent, as observed previously (Figure 2C-D). However, in contrast to transcriptional data, FB3 fibroblasts were marginally depleted in SSc tissues (OR = 0.61). As observed in scRNA-seq analysis, the odds of SSc tissues containing macrophages (OR = 5.98, 95% CI [5.09, 7.08]), FB1 fibroblasts or FB2 fibroblasts (OR = 13.21, 95% CI [10.17, 17.45]) were significantly higher compared with HC tissues **(Fig. 3D**, Table S3).

### Multiome Cells Confirm Cluster Identity Across Modalities

Multiome cells were leveraged to compare clusters between both modalities by projecting cells from the scRNA-seq clusters onto the corresponding cells in the scATAC-seq data. This analysis confirmed that the majority of cells in these clusters matched (Fig. S4-5). Notable differences between the modalities include that FB2 fibroblasts were more transcriptionally diverse than suggested by their complementary epigenomic profiles (Fig. S5), and a subset of these cells cluster with macrophages in scATAC-seq data (**Fig. 2A**, Fig. S3B-C). Additionally, the FB1 cluster of fibroblasts maintains a very distinct epigenomic profile from other fibroblasts, despite having a very similar transcriptomic profile (Fig. S4-5). Integration of epigenetic and transcriptomic data was also accomplished via label transfer based on scATAC-seq predicted gene expression (data not shown). Clusters from both modalities matched in the complementary data type (except for the FB5 cluster) and were placed as expected, based on previous multiome analysis. Although informative, the label transfer was imprecise and didn’t fully recapitulate the clusters. More importantly, label transfer is based on imputed gene expression. Therefore, to avoid diluting the epigenetic data and to ensure the most accurate analysis possible for identifying markers of interest, separately integrated data were used for each modality moving forward.

### Clusters Maintain Distinct Biological Profiles and Highlight Epigenetic Dysregulation

We performed additional analyses to define the biological identity of each of these clusters. To do this, we identified the top differentially expressed (DE) genes (denoted in blue in all figures) and top enriched TF motifs (denoted in red in all figures) for each cluster **(Fig. 4 A-B,** Table S5-6). TF motif enrichment was determined by defining the top differentially accessible peaks (200bp regions of accessible chromatin) and then locating enrichment of TF binding motifs within these peaks. Greater enrichment of a motif within these large accessible chromatin regions is assumed to allow a TF to bind more frequently and approximates TF activity.

**Figure 4.**
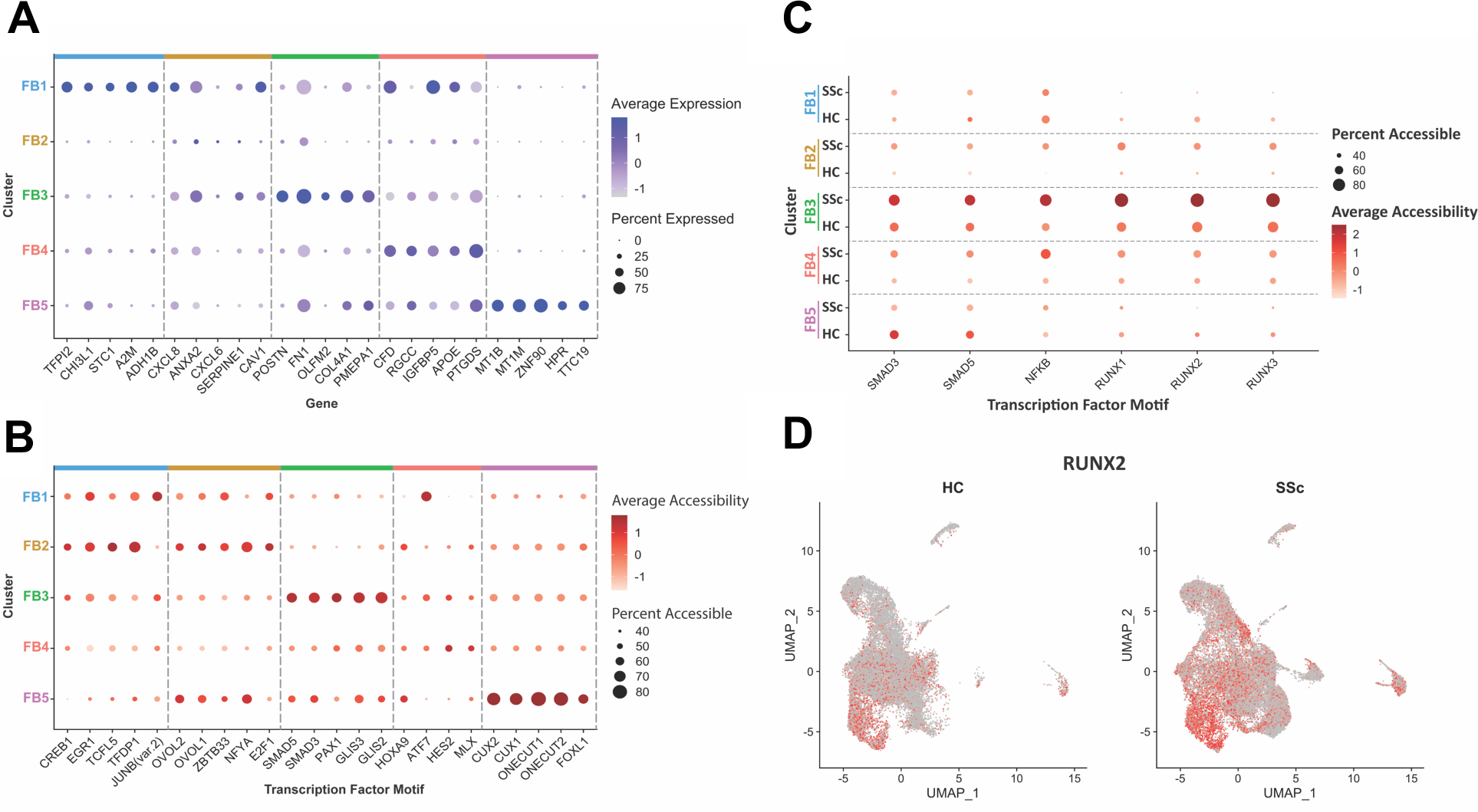
Fibroblast subclusters from saSE tissues are characterized by unique transcriptomic and epigenomic profiles. A) Dot plot featuring top five differentially expressed genes for each saSE tissue fibroblast cluster. B) Dot plot featuring top five differentially accessible transcription factor motifs (JUNB is represented in both FB1 and FB4 clusters). Dot size is representative of percent of cells in this cluster that express the gene or have transcription factor motif accessibility. Average expression or accessibility of cells in each cluster is shown by intensity of color. C) Motif enrichment dot plot of key transcription factors identified in the FB2 fibroblast population split by condition (HC or SSc). D) Feature plot of RUNX2 motif accessibility in HC and SSc saSE tissues. HC = healthy control, SSc = systemic sclerosis

We also investigated the associated biological terms for each cluster using g:Profiler, which predicts gene ontology modules using lists of differentially expressed genes. Key findings are summarized in **Table 1**.

**Table 1:**
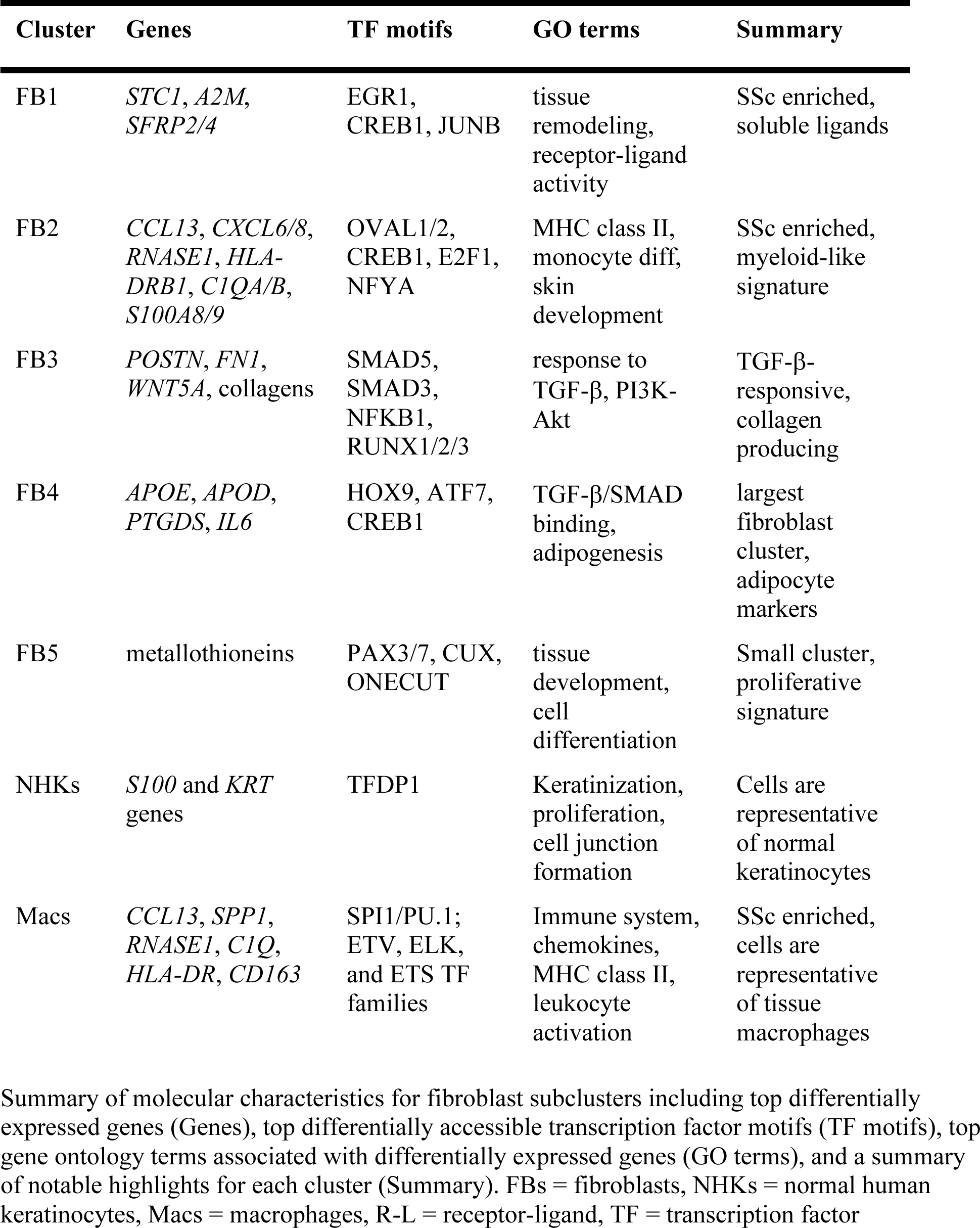
Single-cell multiomic summary of saSE tissue fibroblast subsets.

FB1 was the most significantly enriched cluster in SSc-derived saSE tissues with a high OR in both RNA (OR^RNA^) and ATAC (OR_ATAC_) assays (OR_RNA_ = 9.59, OR_ATAC_ = 13.21). The top DE genes in this cluster included *TFPI2*, *CHI31*, *STC1*, and *A2M* **(Fig 4A**, **Table 1**, Table S5). Notably, *TFPI2* (20), *CHI31* (21), *AGT* (22), and *GDF15* (23) are associated with regulation of TGF-β signaling and fibrosis. Many of these DE genes also code for soluble proteins associated with paracrine signaling. There was also high expression of several genes associated with angiogenesis including CYP1B1, NDUFA4L2, and AGT (24–26). *SFRP2* and *SFRP4* were also enriched in FB1 fibroblasts and have been previously associated with myofibroblasts in single-cell studies of SSc skin (5).

Top enriched TF motifs included CREB1, EGR1, and JUNB **(Fig 4B**, **Table 1**, Table S6). CREB1 regulates *TGFB2* expression (27, 28), and EGR1 and JUNB have been implicated as mediators of fibrosis in SSc (29–34)(35, 36). Enriched gene ontology (GO) pathways included processes related to ECM, tissue remodeling, cell proliferation, angiogenesis, JAK-STAT pathway, TNF signaling, receptor-ligand activity, chemokine/cytokine activity, chemotaxis, and response to hormones and oxygen (Table S7).

FB2 fibroblasts were also enriched in SSc samples (OR_RNA_ = 3.32, OR_ATAC_ = 1.10). Notably, this population was split, with a small portion clustering closely with the macrophages and the other half clustering with fibroblasts **(Fig. 2A**, Fig. S3B-C). Feature plots using QC metrics confirmed that these cells did not have a significant enrichment of doublet cells or ribosomal proteins (Fig. S3D-E). Top FB2 DE genes include many related to immune-regulation, including chemokines (*CXCL8*, *CXCL6*, *CCL13*), cell killing genes (*RNASE1*), MHC class II antigen presentation genes (*CD74*, *HLA-DRB1*), and complement genes (*C1QA*, *C1QB*) **(Fig. 4A**, **Table 1**, Table S5).

Top TFs for this cluster were OVAL1 and OVAL2, which are associated with pluripotency and the specification of cell lineage **(Fig. 4B**, **Table 1**, Table S6) (37). E2F1, which regulates the cell cycle and inflammation (38), and NFYA, which helps orchestrate monocyte differentiation, were also upregulated (39). The CREB1 motif, which was also enriched in this cluster, may play a role in production of TGF-β2 (27). Top GO terms included MHC class II pathways, chemokine and cytokine binding, T-cell activation, monocyte differentiation and pathways associated with ECM deposition, skin development, and angiogenesis (Table S8).

Based on these results and the expression of cell surface markers found on both macrophages (CD14 and CD45) and fibroblasts (COL1A1, THY1), we hypothesize the FB2 cluster is “fibrocyte-like” (40). Fibrocytes (FCs) are thought to be derived from a circulating monocyte pre-cursor population that is capable of trans-differentiating into a fibroblast-like or myofibroblast phenotype. To provide additional insight into the origin of these FB2 cells, we grew self-assembled stromal (SAS) tissues that did not contain monocytes. Integration and clustering analysis indicated that three major fibroblast populations are present in this tissue (Fig. S6A). However, when we projected the FB2 subset score, there was only scattered expression and we were unable to identify a cell population enriched in these cluster-specific genes (Fig. S6B). While the origin and even existence of fibrocytes has long been a topic of debate (40), these findings suggest that regardless of what cell type fibrocytes represent, there are cells characterized by expression of both extracellular matrix and immune genes.

Top DE genes for the FB3 cluster, which was marginally enriched in SSc samples at a transcriptomic level (OR_RNA_ = 1.16, OR_ATAC_ = 0.61), included *POSTN*, *FN1*, *LUM*, and multiple collagens **(Fig 4A**, **Table 1**, Table S5). *POSTN* and *LUM* have been identified as hallmark genes in pro-fibrotic fibroblast populations (41, 42). Additional genes expressed by this population that have been implicated in SSc pathogenesis include collagens, FN1, and *WNT5A* (43, 44).

Top TFs in the FB3 cluster included SMAD5, SMAD3, and NFKB1, which are associated with TGF-β signaling, fibrosis, and inflammation **(Fig 4B**, **Table 1**, Table S6) (45, 46). Other SSc-relevant TFs that showed increased expression in FB3 include TFAP2 proteins (47) and RUNX family proteins, which were identified as enriched regions in our data, with RUNX2 ranking the highest (48). Top GO terms were associated with pathways related to ECM maintenance, response to TGF-β, wound healing, PI3K-Akt signaling pathway, and angiogenesis (Table S9). Visualization of NFKB1, SMAD3, SMAD5, and RUNX family protein motif accessibility confirmed enrichment of accessibility in SSc fibroblasts compared with HC fibroblasts, most notably in FB3 and FB4 fibroblast clusters **(Fig. 4C-D**, Fig. S7A). Gene expression of NFKB1, SMAD3, SMAD5, and RUNX family TFs was also increased in SSc cells across several clusters, primarily in FB1, FB3, and FB4 clusters (Fig. S7B).

*RGCC*, *IGFBP5*, *APOE*, *PTGDS*, *APOD*, and *IL6* were the top DE genes in the largest fibroblast cluster, FB4 (OR_RNA_ = 0.90, OR_ATAC_ = 0.87) **(Fig. 4A**, **Table 1**, Table S5). Each of these genes has been implicated in the pathogenesis of SSc (49)(50). Intriguingly, *RGCC*, *APOD*, and *APOE* are also expressed by adipocytes (51–53); loss of dermal adipose tissue is a hallmark of SSc skin disease (54). TFs most highly represented in this cluster included HOX9, TFE3, ATF7, MYCN, and CREB1, which have also been shown to play roles in adipogenesis and lipid metabolism **(Fig. 4B**, Table S6) (55–58). Notable GO terms included cell migration, angiogenesis, epithelial/smooth muscle cell proliferation, ECM pathways, TGF-β/SMAD binding, and adipogenesis (Table S10).

In contrast to clusters FB1-4, which had greater representation in SSc skin vs. HC skin, FB5 numbers were decreased in SSc skin (OR_RNA_ = 0.59, OR_ATAC_ = 0.45). Several metallothionines (*MT1B*, *MT1M*), which are associated with hyperproliferation of fibroblasts and tissue remodeling, were among the top DE genes in this cluster **(Fig. 4A**, **Table 1**, Table S5) (59). The FB5 cluster was characterized by high levels of PAX3/7, CUX, and ONECUT TFs, which regulate cellular differentiation and chromatin accessibility **(Fig. 4B**, **Table 1**, Table S6) (60)(61)(62). GO terms associated with this cluster were related to tissue formation and maintenance, including tissue development, anatomical morphogenesis, nervous system development, cell differentiation, ECM, and angiogenesis (Table S11). Consistent with DE gene expression results, cell cycle analysis indicated that a large proportion of these cells are proliferative – making it the most proliferative among all fibroblast subpopulations (Fig. S7C).

In accordance with their roles in keratinocyte differentiation and maintenance, the top DE genes in the NHK cluster were *S100* and *KRT* genes (OR_RNA_ = 0.28, OR_ATAC_ = 0.84) (**Table 1**, Table S5) (63, 64). KRT6A and KRT17, which are associated with wound repair and hyperproliferation in psoriasis (65, 66), and S100A8 and S100A9, which have been associated with inflammation (67), were highly enriched. TFs with increased motif accessibility in the NHK cluster included TFDP1, which regulates keratinocyte proliferation (**Table 1**, Table S6) (72), and importantly, the GO terms for this cluster aligned with expected biological processes for keratinocytes. These included terms associated with keratinization, cell proliferation, wound healing, ECM maintenance, and cell junction formation (**Table 1**, Table S12).

*C1Q*, *HLA-DR*, and *CD163*, which are associated with macrophage activation, were among the top DE genes that defined the macrophage cluster (OR_RNA_ = 9.65, OR_ATAC_ = 5.98) (**Table 1**, Table S5) (73) (74). Top TFs included macrophage-specific SPI1/PU.1 (**Table 1**, Table S6) (75), and ETV, ELK, and ETS proteins. Pathways related to the immune system, chemokines, MHC class II molecules, leukocyte activation, and response to stimulus were characteristic of this cluster (**Table 1**, Table S13).

### Fibroblast Subclusters Are Comparable to Clusters Identified in Human Skin

To confirm the presence of these clusters in human skin, we compared results obtained from the HC and SSc fabricated skin model to a single-cell RNA-seq (scRNA-seq) dataset containing both HC donor and SSc skin biopsies (5). Cell types were assigned based on expression of key cell-specific genes identified in skin (Fig. S8), resulting in a fully integrated and clustered scRNA-seq dataset with a total of fourteen clusters of different cell types (Fig. S8A).

The skin data were further clustered to include only fibroblasts, macrophages, and dendritic cells (DCs) (including a small group of JCHAIN expressing macrophages/DCs) (Fig. S9A). This subset of data resulted in a total of five fibroblast clusters and one combined macrophage/DC cluster (Fig. S9B), as validated by cell type-specific markers (Fig. S9C-E). This reference cluster annotation strategy (Fig. S10A) was used to determine which fibroblast clusters in the human skin dataset corresponded to the saSE tissue clusters. This method was validated on the original clustered skin object, and showed high accuracy of placing clusters in this context (Fig. S10B).

Fibroblast subset scores (FB1, FB2, FB3, FB4, and FB5) from saSE tissues were assigned to the re-clustered human skin fibroblasts, confirming an enrichment for these scores in several clusters (**Fig. 5A-C**, Fig. S11A-F). Furthermore, scores for the human skin fibroblast clusters (cluster 0 to cluster 9) matched the cells in this re-clustered dataset, indicating that the original clusters were contained within these newer clusters (**Fig. 5D-E**, Fig. S11G-Q). To validate this assignment, cell names were pulled from the original clustered object and projected onto the new object; this showed similar placement to the cluster scoring strategy (data not shown). Only cluster 1 did not have a clear match but instead corresponded to the FB2 fibroblast population in the saSE data (**Fig. 5D-E**). However, when analyzed separately, cluster 1 showed higher expression of cluster 4 and cluster 0 DE genes (Fig. S11Q). Similar to the FB2 population in the saSE data, cluster 1 did not have high total or unique transcripts (Fig. S3D, Fig. S11R-S), but did contain high mitochondrial gene expression comparable to the macrophage subset (Fig. S11T). Thus, it is possible that this population was filtered out by a more stringent mitochondrial read cutoff in the originally published skin data analysis.

**Figure 5.**
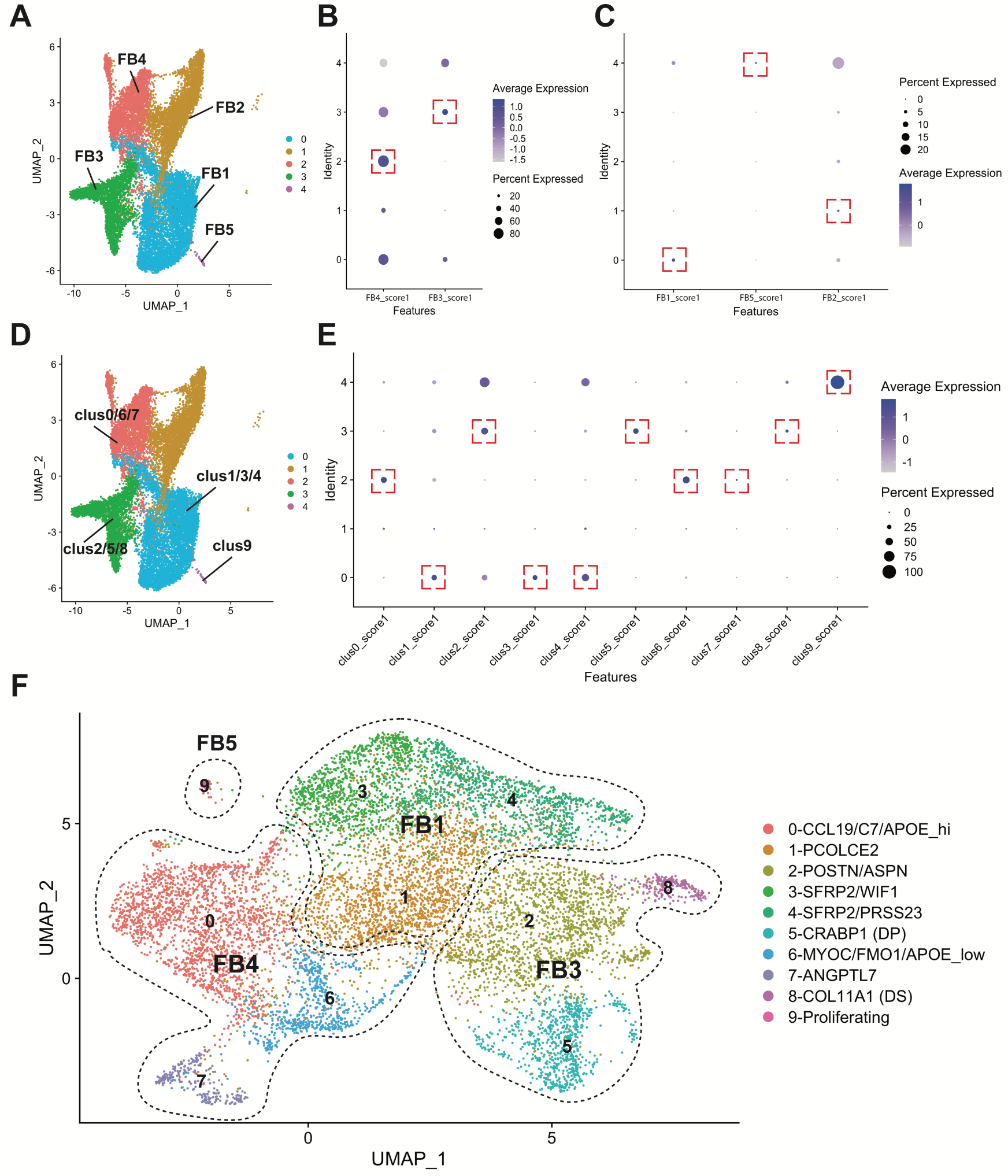
Fibroblast clusters analogous to saSE tissue fibroblasts identified in human skin single-cell RNA-seq dataset. Single-cell RNA-seq data from Tabib *et al.* publication was re-processed using parameters from saSE tissue scRNA-seq data. A) Re-clustered fibroblasts from skin were annotated for saSE tissue fibroblast subclusters by applying a score for the top 50 differentially expressed genes from each cluster and plotting in dot plots (B-C). Cluster scores for fibroblast subclusters FB1, FB2, and FB5 were plotted separately to optimize viewing of these lower expressing genes. C) Similar process was applied to annotate the re-processed and clustered data with the original clusters from the Tabib *et al.* publication (D-E). Annotated UMAP plot (D) and dot plot (E) identified analogous clusters for all except cluster 1 which corresponds to the FB2 cluster of fibroblasts. F) These annotations were transferred to the original clustered fibroblast object from Tabib *et al.* publication for further interpretation of the results.

The original clustered data were manually annotated for ease of interpretation **(Fig. 5F)** using prior analyses **(Fig. 5A-E**, Fig. S11). The FB1 cluster of fibroblasts corresponded to clusters 1, 3 and 4 in the skin data. Consistent with the FB1 cluster, fibroblast clusters 1, 3 and 4 expressed *SFRP2* and *SFRP4* transcripts and were similarly identified in the comparator human skin dataset as enriched in SSc samples, with predicted pathogenic differentiation into the terminal cluster 4 fibroblast population (5). The FB3 population matched fibroblast clusters 5 and 8 in the comparator skin data, and are collagen-producing, consistent with the high ECM expression profile of the FB3 fibroblasts. The FB4 cluster approximated fibroblast clusters 0, 6, and 7 in human skin, and were characterized by *APOE* and *APOD* expression. Cluster 9 in skin data set matched the FB5 population; both were characterized by cell proliferation. Thus, manual comparison of highly expressed genes in these clusters provided additional validation for the unbiased assignment of analogous clusters across these two datasets.

### Fibroblast Epigenetic Dysregulation Highlights CREB1/EGR1 Axis in FB1 Enrichment

To better understand the epigenetic mechanisms driving differential cell type enrichment and pathological tissue phenotype, we compared the epigenetic profile of SSc cells from each cluster to their HC counterparts. Global analysis using all cells in the tissue identified TFs from enriched clusters (such as OVOL2, ELK1, CREB1, and EGR1) (Table S14). However, other TFs independent of cluster enrichment also demonstrated increased motif accessibility in SSc. Among these TFs, HIFNP, HIF1A, and XBP1 ranked among the highest differentially accessible motifs as well as SMAD3 and RUNX2.

A similar analysis was performed to identify differentially accessible TF binding motifs between disease states for each independent cluster. Only three cell clusters had statistically significant differentially accessible TF motifs: FB2, FB3, and FB4 (Table S14). Notably, these three clusters shared TF motifs (Table S15), intersecting in EGR1, which was observed to have high motif accessibility in the FB1 SSc-specific subset **(Fig. 4B**, Table S6).

Dot plots visually demonstrate an increase in EGR1 motif accessibility and percent of cells with accessibility in this motif in all SSc fibroblast clusters **(Fig. 6A**, Fig. S12A). Notably, differential gene expression of these TFs was minimal, with the most pronounced enrichment of the HIF1A motif across FB4, FB3, and FB1 clusters (Fig. S12B). We hypothesized that if EGR1 motif accessibility were increased in SSc fibroblasts across the clusters, this may drive differentiation of a pathogenic fibroblast population characterized by increased EGR1 accessibility (i.e., the FB1 cluster enriched in SSc samples). To test this, a subset of fibroblasts (FB1, FB3, and FB4 excluding the FB2 population) from the integrated scATAC-seq data as described previously using clustree was isolated and re-clustered. At this resolution, two cell populations were identified as enriched in SSc tissues **(Fig. 6B)** – the FB1 cluster and a population that arises from a subset of the FB4 cluster (Fig. S13A-B), designated as the FB4.2 cluster. Notably, FB4.2 was distinctly clustered in scATAC-seq data, but was dispersed across the FB4 population in transcriptomic data (Fig. S13C-D).

**Figure 6:**
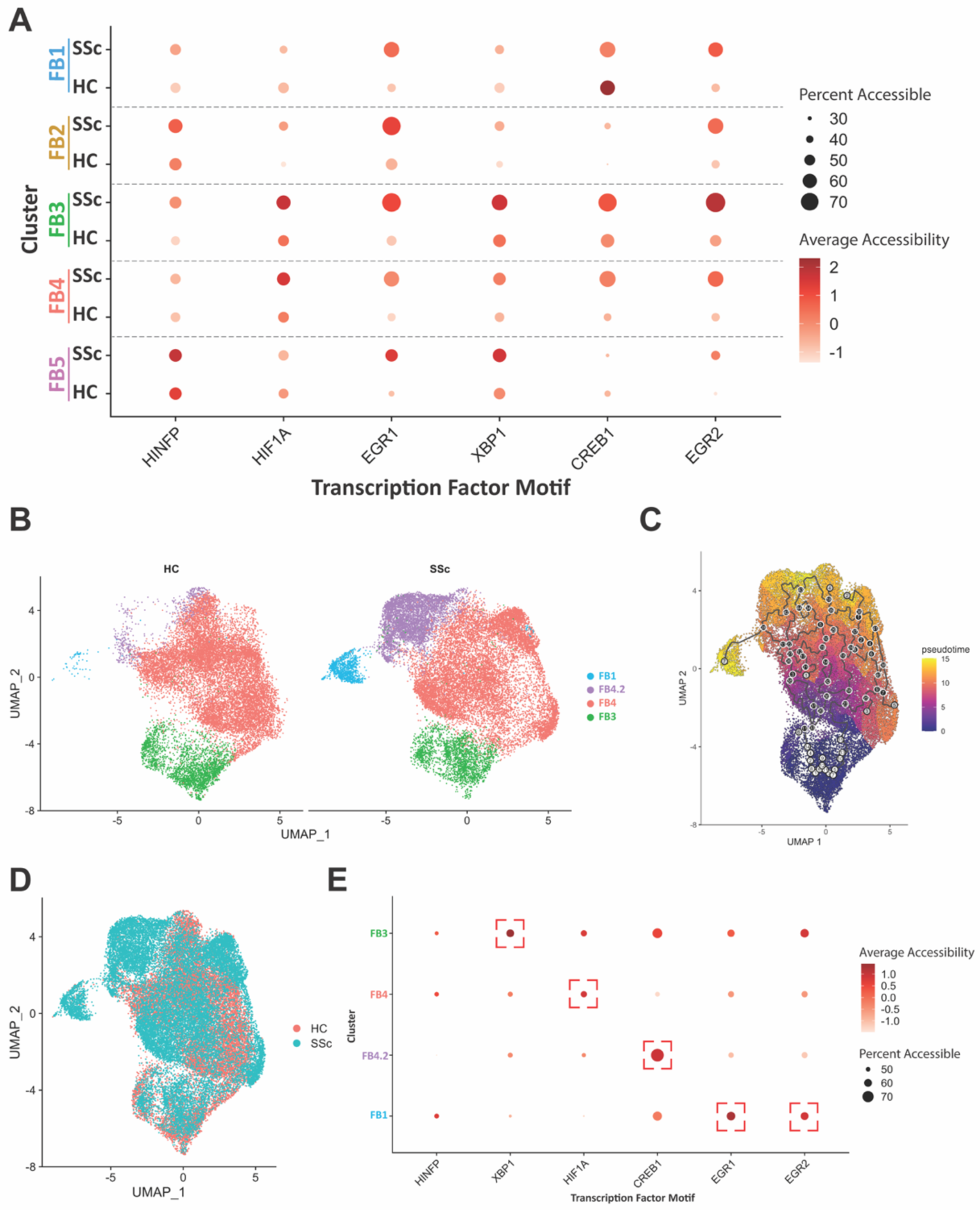
Pseudotime analysis predicts FB1 fibroblasts arise from FB3 fibroblasts via transcription factor regulatory cascade. A) Dot plot of transcription factor motif accessibility across SSc-enriched TF motifs split by condition (HC or SSc). B) UMAP projection of re-clustered single-cell ATAC-seq (clusters FB1, FB3, and FB4) split by HC and SSc samples to show disease-specific enrichment of FB1.2 and FB3 clusters. C) Pseudotime analysis showing projected cell differentiation trajectory with FB3 cells as the starting population. D) Same clustered object with cells colored by condition (HC or SSc) to highlight global differences in epigenomic profiles. E) Dot plot of transcription factor motif enrichment for motifs determined to be enriched in SSc fibroblast subsets. Size of dot corresponds to percent of cells with accessible motif in that cluster and intensity of color corresponds to average accessibility of that motif. HC = healthy control, SSc = systemic sclerosis, saSE = Self-Assembled Skin Equivalent

Genes enriched in the FB4.2 cluster include *KCND2*, *C7*, *PDK4*, *BMPER*, *EREG*, *PLOD2*, and *PAX3* (Table S5), which are involved in the regulation of wound healing and fibrosis (76–81). There was overlap with the FB4 cluster among a subset of genes, including *IL6*, *APOE*, and *VEGFA*.

Differentially accessible transcription factor motifs in the FB4.2 cluster include CREB1, XBP1, JUN, and HIF1A (Table S6). GO terms enriched in FB4.2 fibroblasts are primarily related to pathways of tissue morphogenesis, angiogenesis, epithelial cells signaling and migration, ECM, and cell junctions (Table S16), suggesting a potential role for this cell population in the pathogenesis of SSc.

FB3 cells were defined as the root population based on the TGFB1-responsive molecular phenotype and a reverse trajectory starting with the FB1 population (Fig. S13E-F). With FB3 cells as the root population for differentiation of FB1 cells, a pseudotime trajectory was developed that moved from FB3 cells along the outer edge of the FB4 population, to the FB4.2 cluster, and finally to FB1 populations **(Fig. 6C)**. Of note, this route followed areas of the FB4 cluster with a high density of SSc fibroblasts, indicating that these cells are already epigenetically shifted towards a pathogenic profile **(Fig. 6D).** Visualization of the top SSc-specific dysregulated motifs in each of these specific fibroblast clusters showed enrichment in SSc tissues, and the FB3 population contained the clearest enrichment of accessibility in all motifs **(Fig. 6A, E**; Fig. S12A). FB4 was more skewed towards HINFP and HIF1A motif enrichment; FB4.2 was most clearly enriched in CREB1 motifs; and FB1 was clearly most enriched in EGR1, with increased accessibility of CREB1 and EGR2 motifs (**Fig. 6E**, Fig. S12).

In further support of this, analysis of the human skin scRNA-seq data set indicated dysregulation of *EGR1* in SSc fibroblast and macrophage populations (Fig. S14) (5), and *EGR1* expression was similarly increased in SSc fibroblasts (primarily FB1-like, FB3-like, and FB4-like) and SSc macrophages/DCs (Fig. S14C).

### SSc Macrophages Are Characterized by an Activated Epigenetic Profile

Macrophages were significantly enriched in SSc-derived saSE tissues compared with HC tissues **(Fig. 2-3)** and have been implicated as important mediators in SSc fibrosis (82). Prior work has shown that human SSc macrophages have a pro-fibrotic immunophenotype, and both *in vivo* and *in vitro* model systems support a role for macrophages in the induction of fibroblast activation in SSc (83). Consistent with prior observations in human skin (84), SSc saSE tissues contain greater numbers of CD163^+^ and CD206^+^ macrophages than HC tissues (8), which is also reflected in increased CD163 and CD206 surface marker levels in SSc tissues (Fig. S15). Flow cytometry analyses also confirmed that macrophages comprised ∼5-10% of cells in the saSE tissue, which is consistent with reported cell proportions in human skin (6, 85).

To define the epigenetic state of macrophages in saSE SSc and HC tissues, macrophages were isolated and re-clustered (**Fig. 7A**, Fig. S16). Clustree analysis identified three distinct epigenetic macrophage populations, which were named Macs_A, B, and C (**Fig. 7A**, Fig. S16A-B). Because LPS, INFγ, and IL-4 have been implicated in SSc pathogenesis (86–88), relative levels of TFs associated with LPS/INFγ (IRF3, IRF5, IRF7, IRF9, STAT1, and p65) vs. IL-4 (IRF4, STAT3, PPARG, KLF4, HIF1A, p50/NFKB1) activation were compared in the clusters. This was used to generate “LPS/INFγ” and “IL-4” scores which were applied to determine enrichment of these signatures in macrophage clusters (**Fig. 7B**) (89–92). Both LPS/INFγ and IL-4 scores were enriched in SSc tissues (Fig. S16C). Consistent with prior published data indicating human SSc macrophages are characterized by both pro-inflammatory and pro-fibrotic activation, there was highly significant overlap in most cells of the LPS/INFγ and IL-4 TF score (**Fig. 7C**). Fold enrichment and odds ratio analysis showed that the Macs_B cluster was the most significantly enriched in SSc samples (**Fig. 7D-E**).

**Figure 7.**
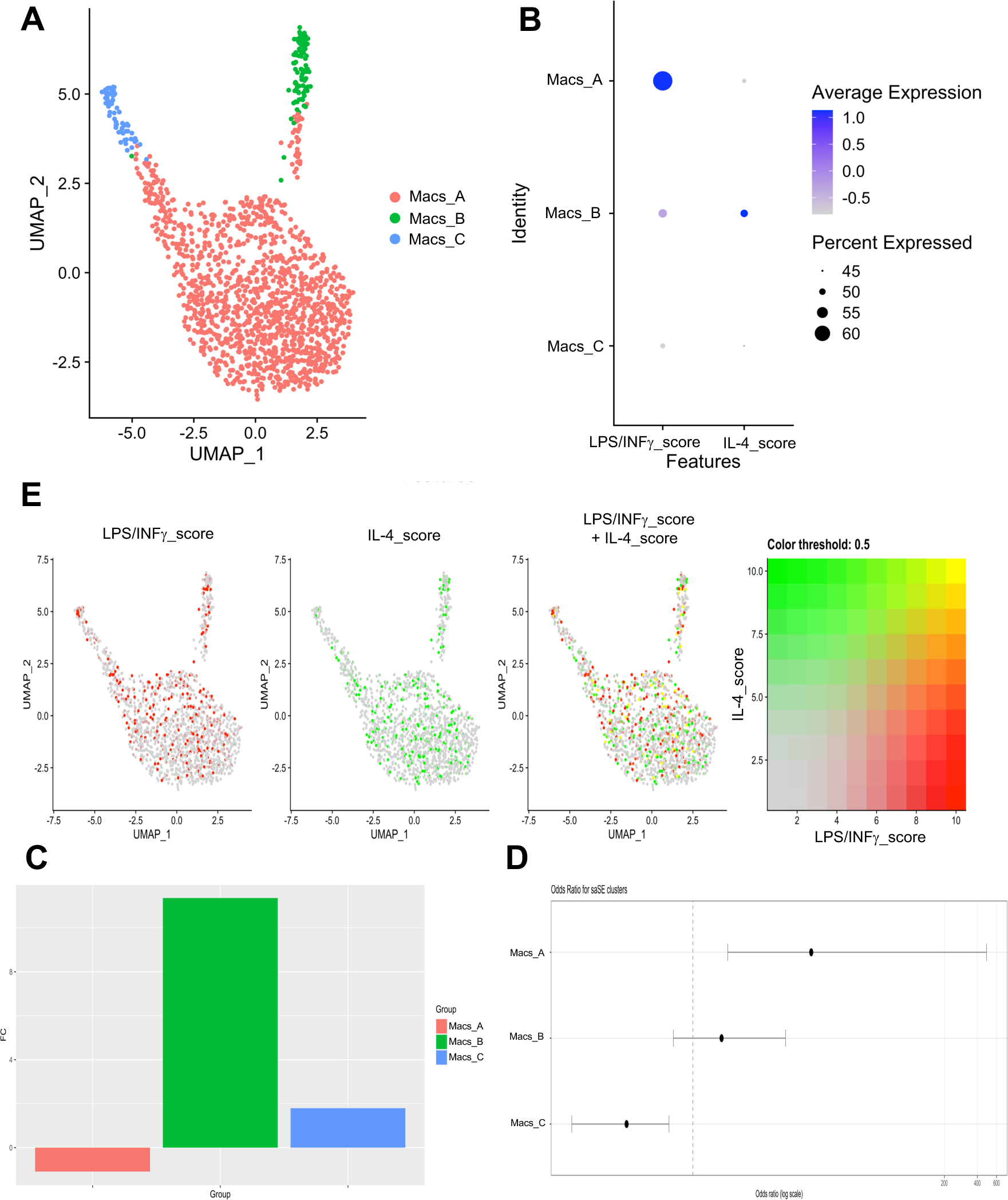
Activated LPS/INFγ and IL-4 profiles of macrophages from systemic sclerosis (SSc) saSE tissues is demonstrated using single-cell ATAC-seq data. A) UMAP projection of clustered macrophages with subclusters determined. B) Dot plot of LPS**/**INFγ and IL-4 signatures for each macrophage sub-cluster based on transcription factor motif enrichment score for each cell. C) LPS**/**INFγ, IL-4, and overlapping scores projected onto macrophages using UMAP. D) Fold change in proportion of macrophages in each cluster in SSc vs. healthy control (HC) tissues. E) Odds ratio of cells present in each macrophage cluster in SSc vs. HC. HC = healthy control, SSc = systemic sclerosis, saSE = Self-Assembled Skin Equivalent, Macs = macrophages.

### Ligand-Receptor Analysis Implicates FB1-derived Ligands in Macrophage Activation

Because FB1 cells differentially express many soluble ligands that mediate cell activation (**Table 1**, **Fig. 4A**, Table S5), we hypothesized that FB1 cells may engage with macrophages to regulate fibrosis in SSc via ligand-receptor (L-R) interactions. To address this, the nichenetr package was used to predict L-R interactions **(Fig. 8)**. NicheNet analysis predicts active L-R interactions based on correlation between ligands in the sender cell type and target genes of interest in the receiver cell type **(Fig. 8E**, Fig. S17A) (93). By identifying ligands that are predicted to modify the gene expression of target genes in receiver cells, the results are focused on the interactions most likely to impact overall biology.

**Figure 8.**
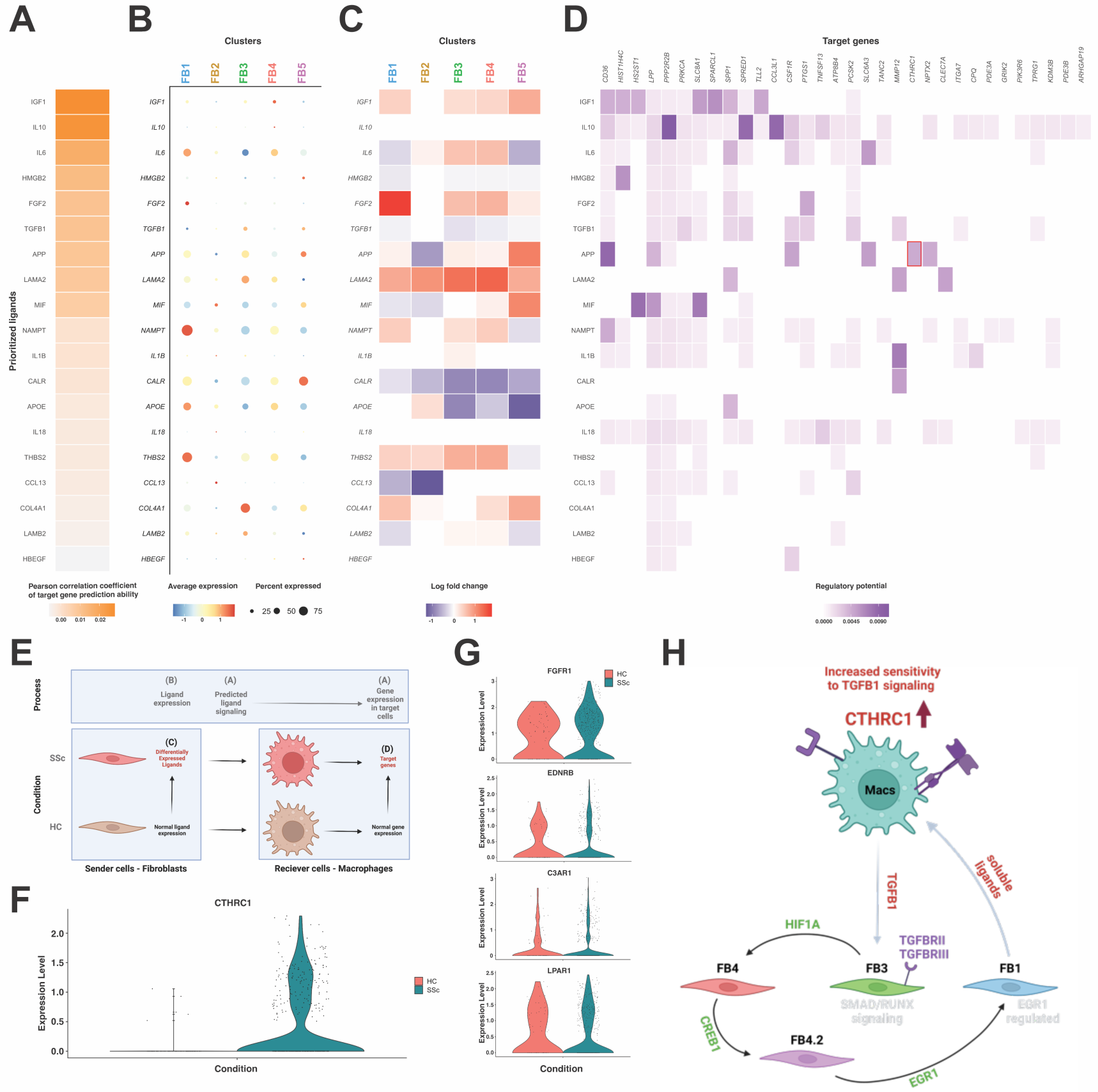
Ligand-receptor analysis indicates fibroblast-macrophage crosstalk results in increased expression of CTHRC1. A-D) Ligand-receptor NicheNet analysis of scRNA-seq and data from saSE tissues. E) Schematic of NicheNet analysis. Fibroblast subsets (FB1, FB2, FB3, FB4, and FB5) are defined as senders and macrophages as receivers. Target genes in macrophages are defined as differentially expressed genes in SSc macrophages from saSE tissues. A) Top ranked ligands based on correlated co-expression of ligand expression in fibroblasts and target genes expression in macrophages. B) Gene expression of predicted ligands in fibroblast clusters. C) Log fold change of predicted ligands in SSc fibroblasts from each subcluster. D) Target genes in macrophages predicted to be regulated by ligands from fibroblast clusters. F) Violin plot of CTHRC1 gene expression from saSE scRNA-seq data split by condition (HC or SSc). G) Violin plot of FGF2 receptor (FGR1) and APP receptor (EDNRB, C3AR1, and LPAR1) gene expression from saSE scRNA-seq data split by condition (HC or SSc). J) Schematic representation of predicted saSE fibrosis feedback loop summarizing potential interactions between fibroblasts and macrophages in SSc tissues. Black arrows indicate cellular differentiation. Transcription factors in green are predicted to mediate fibroblast differentiation. Grey arrows indicate ligand-receptor interaction with ligand genes/proteins in red. Cell surface receptors are denoted in purple. HC = healthy control, SSc = systemic sclerosis, FB = fibroblast, saSE = self-assembled skin equivalent

Given our interest in understanding the potential influence of soluble ligands produced by FB1 cells on myeloid activation, all fibroblast subsets in our tissue model were defined as senders and macrophages as receivers. Results of this analysis showed that the FB1 population was enriched in nearly all the top ranked ligands predicted to interact with macrophages, and had multiple DE ligands based on the disease state of the cells **(Fig. 8A-C)**. Of these ligands, *FGF2* was the most robustly expressed in FB1 cells, with a high fold-increase in disease state **(Fig. 8B, C)**, and the cognate FGF2 receptor *FGR1* was increased in SSc macrophages **(Fig. 8G**, Fig. S17C). *APP* was also differentially expressed in multiple fibroblast subsets. **(Fig. 8B, C).** One of the predicted target genes upregulated in macrophages was *CTHRC1* **(Fig. 8F**, Fig. S17B), which regulates TGF-β signaling and was predicted to be upregulated by APP **(Fig. 8D)**. Notably, expression of the APP receptors *C3AR1*, *EDNRB*, and *LPAR1* was elevated in SSc macrophages **(Fig. 8G**, Fig. S17D). Based on these collective results, we developed a model of cooperative interaction between fibroblasts and macrophages that orchestrates inflammation and fibrosis in SSc (**Fig. 8H**).

## Discussion

This study identifies and validates the presence of key pathogenic fibroblast subpopulations using a disease-specific *in vitro* model of SSc skin-fibrosis and defines the epigenetic dysregulation of these populations at single-cell resolution using a multi-omic approach. Tissues fabricated from patient-derived cells and plasma displayed histological evidence of a pro-fibrotic phenotype. These measures were consistent with results from Huang et al. that demonstrate the saSE tissue model recapitulates the phenotypic and molecular dysregulation observed in SSc skin biopsies (8). In addition, multimodal single-cell analysis identified macrophages, keratinocytes, and five major fibroblast subsets that were found in both HC and SSc tissues. The fibroblast heterogeneity is notable and underscores the importance of findings from a recent study by Kosarek *et al.* showing that the tissue microenvironment created by the saSE model is able to capture fibroblast heterogeneity that is lost in conventional fibroblast monolayer cultures (94). This demonstrates the critical importance of studying SSc pathogenesis using *in vitro* systems that most closely replicate the complexity of SSc in human tissues.

Since the same number of cells was seeded for all tissues, subtype-specific fold enrichment of these clusters could be accurately calculated. This analysis revealed that SSc-derived tissues were more highly enriched in fibroblast FB1 and FB2 populations and macrophages compared to HC-derived tissues. Molecular profiling for each of these populations defines broad characteristics that define the various fibroblast subtypes. The FB1 population was the most enriched in the SSc-tissues but was characterized by transcripts of soluble signaling molecules (A2M, TFPI2, STC1) as opposed to ECM-related proteins. STC1 is a secreted factor that in the context of cancer has been shown to modulate macrophage activation (95). A2M is a secreted plasma protein that binds cytokines and growth factor proteins, readily enters macrophages with cargo to modify cellular phenotype, and has been shown to promote antigen presentation for major histocompatibility complex (MHC)-mediated T-cell activation (96). Furthermore, this cluster differentially expressed *SFRP4*, suggesting that it might be molecularly similar to the myofibroblast SSc-specific cluster previously identified in single-cell transcriptional data from human skin (5). Top differentially accessible TF motifs for this cluster implicated EGR1 as important in regulating this unique pathogenic epigenetic state.

A novel finding from this study is identification of the FB2 cluster of fibroblasts, which was moderately enriched in SSc samples and expressed many immune-related genes, TFs, and pathways. Our analysis suggested that FB2 cells may represent fibrocytes (40). Circulating fibrocytes are enriched in the blood of SSc patients and have been shown to correlate positively with mRSS (100) and severity of lung fibrosis in SSc patients (101–103). In published scRNA-seq data from human skin (5), this cluster of cells did not match any of the gene expression cluster scores for fibroblast clusters previously reported for this dataset, but did express higher levels of mitochondrial genes, suggesting this cluster may have been filtered out in previous analyses. Comparable cell types from human tissues have also been described in recent single-cell literature (40). Considering the lack of clarity about the presence and function of fibrocytes in various disease states (40) our findings of this putative fibrocyte cluster suggest the participation of fibrocytes in fibrotic disease states as a tissue-based phenomenon. As a result, further rigorous studies are warranted to fully determine the heterogeneity of this cluster and potential precursor populations and to determine what role these cells play in fibrosis and SSc pathogenesis. Additional experiments should include lineage tracing or detailed time course studies and further *in vitro* functional characterization of cell subsets within this cluster.

In contrast, the FB3 population was not consistently enriched in SSc tissues but was characterized by a heightened TGF-β-responsive TF motif profile (SMAD3, SMAD5, NFKB1, and RUNX proteins), and this profile was even more pronounced in SSc tissues. In addition, the FB3 population expressed multiple collagens and pro-fibrotic genes (*POSTN*, *LUM*) previously associated with fibrosis in other models (41, 42). Collectively, these data suggest that although FB3 is not significantly enriched in SSc samples, it still maintains a pro-fibrogenic profile that is differentially regulated based on disease state, with an increase of TGF-β-related TFs. The FB4 population was characterized by several adipocyte-associated molecules and GO pathways, suggesting it may be related to adipocyte-derived fibroblasts (54, 97). Of note, *IL6* expression was elevated in this cluster, implicating a role for stromal cells as contributors to high IL6 levels in SSc skin, sera, and supernatants of SSc-derived tissues (8, 50). The FB5 population represented a rare hyperproliferative population of fibroblasts with potential for cellular renewal, as indicated by the differential accessibility of stem cell-like TFs PAX3/7.

Comparable fibroblast subpopulations were identified in human skin data using DE gene scores for each cluster, suggesting these populations approximate those observed *in vivo.* These findings also reflected the presence of both papillary and reticular fibroblast populations as previously identified and described in human skin based on work by Watts and Lafyatis (49, 98, 99).

Analysis of disease-specific epigenetic changes in specific fibroblast populations identified differential accessibility of HINFP, HIF1A, CREB1, and EGR1 motifs in several fibroblast clusters. Pseudotime analysis suggested that FB3 fibroblasts may give rise to FB1 fibroblasts via a TF cascade. This cascade started with increased HINFP activity in FB3 cells, then HIF1A activity in FB4 cells, CREB1 activity in an SSc-enriched subset of FB4 cells (FB4.2), and finally EGR1 TF activity in the FB1 cells. Because CREB1 regulates expression of EGR1 (104), a model in which dysregulation by EGR1 is preceded by CREB1 dysregulation is logical and suggests that pathogenic differentiation may occur prior to EGR1, even though EGR1 characterized the terminal differentiated state.

EGR1 has been associated with SSc in multiple studies from patient tissues, including a recent bulk ATAC-seq study of SSc fibroblasts and epithelial cells, which concluded that EGR1 was one of the key dysregulated TFs (29–32, 34). In addition, several studies have shown the efficacy of EGR1-targeted therapies in reducing fibrosis in both preclinical models and in clinical trials for rheumatoid arthritis (31, 105, 106). Given the evidence from this and other studies, it is possible that an anti-EGR1 therapeutic strategy may have the potential to treat fibrosis in SSc skin.

Macrophages were highly enriched in SSc compared with healthy control tissues, which is consistent with observations in human skin biopsies from patient samples (9). Because the same number of macrophages was added to each tissue, we hypothesize that SSc tissue macrophages persist due to sustained activation derived, at least in part, from fibroblast-secreted factors. Ligand-receptor interaction studies support this model (Figure 8).

Finally, analysis of single-cell data revealed that keratinocytes in the 3D tissue model express keratin subtypes such as KRT6A, KRT16, and KRT17, which are associated with hyperproliferation and elevated wound healing (64–66). This demonstrates that 3D skin tissues may be used to further investigate the contribution of epidermal-mediated crosstalk to the pathophysiology of SSc. It is well-established that SSc keratinocytes activate pro-fibrotic phenotypes in skin that contribute to SSc (107–109); however, the impact of epidermal keratinocyte-derived growth factors in tissues and how they activate dermal fibroblasts to drive fibrosis is not yet fully understood.

In this study, we have elucidated, for the first time, epigenetic regulation of SSc fibroblasts and macrophages using paired scRNA-seq and scATAC-seq in a novel 3D fabricated skin model that accurately recapitulates disease. These studies identified specific fibroblast subsets and pathways of epigenetic dysregulation that are enriched in SSc tissue, providing insight into the underlying mechanisms of pathogenesis that underpin this disease. Consistent with observations in patient skin, macrophage numbers were increased in this model, and L-R analysis identified candidate regulators of SSc fibroblast/macrophage activation. Collectively, these results demonstrate the need for a tissue system like the saSE model to more closely mimic the fibrotic tissue microenvironment for investigation and development of novel, targeted, much-needed treatments for SSc at single-cell resolution.

## Materials and Methods

### Patients

Patients were recruited as described (8). Demographic and clinical donor characteristics for primary cells and plasma used to construct tissue were outlined in **Supplemental Table S1**.

### Isolation of Human Cells and Plasma

Single-cell analyses were performed on saSE tissues cultured from autologous SSc patient or HC dermal fibroblasts, monocytes, plasma and NHKs as described (8). Whole blood samples were treated with heparin, and cellular components were separated from plasma supernatant by centrifugation at 400g for 35 minutes at 20°C. Peripheral blood mononuclear cells were isolated from heparinized paired donor blood using Ficoll-Paque Premium (density 1.077 g/ml). Monocytes were purified from mononuclear cells using CD14^+^ microbeads beads (Miltenyi). This method of isolation routinely generates monocytes that are >98% pure. Monocytes and plasma were stored at −80°C and thawed once prior to use.

Fibroblasts were isolated as previously described (110). Briefly, a 4mm skin biopsy was obtained from the forearm/bicep of SSc patients and healthy control donors. The epidermis was removed by overnight incubation in Dispase (Invitrogen) at 4°C, and then samples were minced and incubated while stirring for 1hr at 37°C in Dulbecco’s modified Eagle’s medium/F-12 (Invitrogen) media containing collagenase (Invitrogen) and hyaluronidase (Sigma-Aldrich). Cells were collected by centrifugation and cultured in DMEM with 10% Hyclone fetal bovine serum (FBS).

### 3D Tissue Culture

Self-assembled skin-equivalent (saSE) tissues were cultured as previously described (8). Briefly, fibroblasts and monocytes were cultured in Transwells for two weeks prior to feeding with plasma-conditioned media. At week three, keratinocytes were layered on top of the dermal layer. After a week, a liquid-air interface was raised for cornification of the epithelial layer. Tissues were harvested at five weeks for histology and bulk RNA, or subjected to tissue digestion for 10x protocols. Self-assembled stromal (SAS) tissues containing no monocytes or NHKs were grown from dermal fibroblasts and fed with plasma-supplemented media at two weeks as previously described (16).

### Histology

Transwell membranes containing saSE tissues were harvested and placed in histology cassettes with foam inserts. The tissues were fixed overnight in 10% formalin and paraffin embedded. Tissues were sliced parallel to the cassette to create a 4μm cross section of the center of the tissue prior to H&E or immunohistochemistry (IHC) staining with anti-αSMA antibody. For H&E staining, sections were dried at RT before loading on the Tissue-Tek Prisma Stainer (Sakura Finetek USA) and running the automated routine H&E program. IHC slides were air dried before baking at 60°C for 30 min. The Leica Bond Rx (Leica Biosystems Inc.) automated system was run, including paraffin dewax, antigen retrieval, peroxide block, and staining. Heat induced epitope retrieval was achieved via incubation of the slide at 100°C for 20 min in Bond Epitope Retrieval 1, pH6.0 (Leica Biosystems AR9961). Slides were then incubated with 1:200 primary antibody (anti-alpha smooth muscle actin (αSMA), Abcam ab5694) for 15 min at RT. Primary antibody binding was detected and visualized using the Leica Bond Polymer Refine Detection Kit (Leica Biosystems DS9800) with DAB chromogen and Hematoxylin counterstain. Histology metrics were obtained using the opensource software QuPath (version 0.2.3). Tissue thickness was calculated using the average of three individual measurements at roughly equal intervals across the length of the tissue. QuPath software was used to detect αSMA-producing cells and positive cells were reported as a percentage of the total detected cells in the tissue (using hematoxylin staining of the nuclei). Tissue areas were determined using overhead images and QuPath software.

### Tissue Digestion

Tissues were minced and digested in 1 ml collagenase digestion buffer (1 mg/ml Collagenase II and 2 mg/ml DNase I in RPMI) at 32°C for 40-50 min while stirring. Digestion was quenched by addition of flow buffer (20 mM EDTA and 0.2% w/v BSA in PBS). Tissue digestion was strained through 70 μm filter, pelleted at 1800 rpm for 8 min, resuspended in 100 μl flow buffer, and strained through 40 μm FlowMi® filter tip. Final single-cell suspension was subjected to flow cytometry and 10x single-cell/multiome protocols.

### Flow Cytometry

Tissues constructed for flow cytometry analysis were non-autologous and used healthy control monocytes with disease-matched fibroblasts and plasma (donor details in Table S4.1), which has been shown to induce the SSc macrophage phenotype (Bhandari 2020). Additional details on fluorophore-conjugated antibodies can be found in **Supplemental Table S18**. Cell staining was performed for one hour at 4C with 2 mg/ml Globulins Cohn fraction II, III block buffer (Sigma) to reduce non-specific FcR binding. In all conditions, doublets were excluded by forward scatter A (FSC-A) versus FSC-H gating. Cells were analyzed using an 8-color MACSQuant 10 (Miltenyi Biotec) with 3 laser sources (405 nm, 488 nm, 635 nm) and FlowLogic 501.2A software (Inivai Technologies).

### Single-cell and Multiome Sequencing

Several saSE tissues were processed and pooled for each single-cell sample to capture technical variability and ensure a high enough cell number for sequencing. This required a total of 41 individual 3D tissues (HC1 n=6, HC2 n=7, HC3 = 8, SSc1 n = 8, SSc2 n=6, SSc3 n=6). Tissue samples for HC2, HC3, and SSc1 were digested, cells pooled, and split into two separate samples, which were subjected to 10x 3’ scRNA-seq v3.1 and scATAC-seq v1.1 protocols, respectively. Each sequenced sample contained approximately 5,000 cells with a sequencing depth of 12,000-18,000 reads per cell. Tissue samples for HC1, SSc2, and SSc3 were digested and subjected to 10x multiome protocol. Each sequenced sample contained approximately 11,000-13,000 cells per sample with a sequencing depth of 4,000-5,000 reads per cell. SAS tissues were digested and processed for 10x 3’ scRNA-seq v3.1 with approximately 14,000-22, 000 cells per sample with a sequencing depth of 700 reads per cell. ATAC-seq data and scRNA-seq data were pre-processed using v2 and v1 Cell Ranger, respectively.

### Statistical Analyses

Briefly, differential gene expression or accessibility was determined using nonparametric Wilcoxon Rank Sum test. Proportional differences in cell clusters were determined using log fold change or odds ratio analyses. Statistical significance was determined using Students T-test or Wilcoxon Mann-Whitney tests as appropriate and a p-value of <0.05 was considered significant. All data were analyzed using R (v4.2.0). Single-cell data were analyzed using Seurat v3, and Signac 1.7.0 packages (111–117). The clustree package was used to determine cluster resolution (118). Packages chromVAR and JASPAR were used to annotate transcription factor motifs (119, 120). The online tool g:Profiler was used for gene ontology (GO) term analysis (121, 122). Monocle3 was used for pseudotime analyses and nichenetr for ligand-receptor analyses (93, 123). More detailed computational and statistical methods can be found in the Supplemental Methods.

### Study Approval

The study was approved by the Committee for the Protection of Human Subjects at Dartmouth College (STUDY00024329: *Autoimmune Disease Pathogenesis and Treatment via Integrative Genomics*) and the Institutional Review Board at Tufts (12773: *Complex 3-Dimensional in Vitro Human Skin Models for Scleroderma*, 01535: *Functional analysis of cellular diversity and cell-cell interactions in Scleroderma 3D skin-like tissues*). All participants provided written informed consent.

## Data Availability

Raw files for single-cell data are posted online in the Gene Expression Omnibus (#TBD). Other data and R analysis files used in this study are available from authors upon request.

## Author Contributions

TRA drafted the manuscript with contributions from all authors. All authors edited and approved the final manuscript for publication. TRA, NNK, MLW, PAP, and JG designed the study. MH, AS, and DT aided in experimental design and protocol optimization. TRA, NNK, RP, HJ, HY, and TW contributed to tissue culture, histology, and processing of single-cell samples. TRA, NNK, MH, DT, GT, and RB contributed to flow cytometry tissues and processing. TRA performed all statistical analysis and data visualization with support from NNK, MM, and DP.

## Acknowledgements

This work was supported by grants from the National Institute of Arthritis and Musculoskeletal and Skin Diseases (NIAMS) R43 and R44 AR072170 (MLW and JG), NIAID R21AI169420 (PAP and MLW) and NIAMS R21AI178651 (PAP and MLW), the Scleroderma Research Foundation (MLW), and the Burroughs-Wellcome Big Data in the Life Sciences Fellowship (TRA). Studies utilized the Dartmouth Center for Quantitative Biology (CQB) Single Cell Genomics (SCG) and Genomic Data Science (GDS) cores supported by National Institutes of Health (NIH) National Institute of General Medical Sciences (NIGMS) Centers of Biomedical Research Excellence Grant P20GM130454. We acknowledge the Clinical Genomics and Advanced Technology (CGAT) laboratory in the Department of Pathology and Laboratory Medicine of the Dartmouth Hitchcock Health System, the Immune Monitoring Lab, and the Pathology Shared Resource at the Dartmouth Cancer Center with NCI Cancer Center Support Grant 5P30 CA023108-37.

We acknowledge Dr. Robert Lafyatis for providing pre-analyzed data for validation of the cross-dataset single-cell cluster annotation strategy. We thank Dr. Fred Kolling for consultation, processing, and sequencing of 10x single-cell and single-nucleotide samples.

