## Supplemental Figures for "Single-cell epigenomic dysregulation of Systemic Sclerosis fibroblasts via CREB1/EGR1 axis in self-assembled human skin equivalents"

### Supplemental Materials

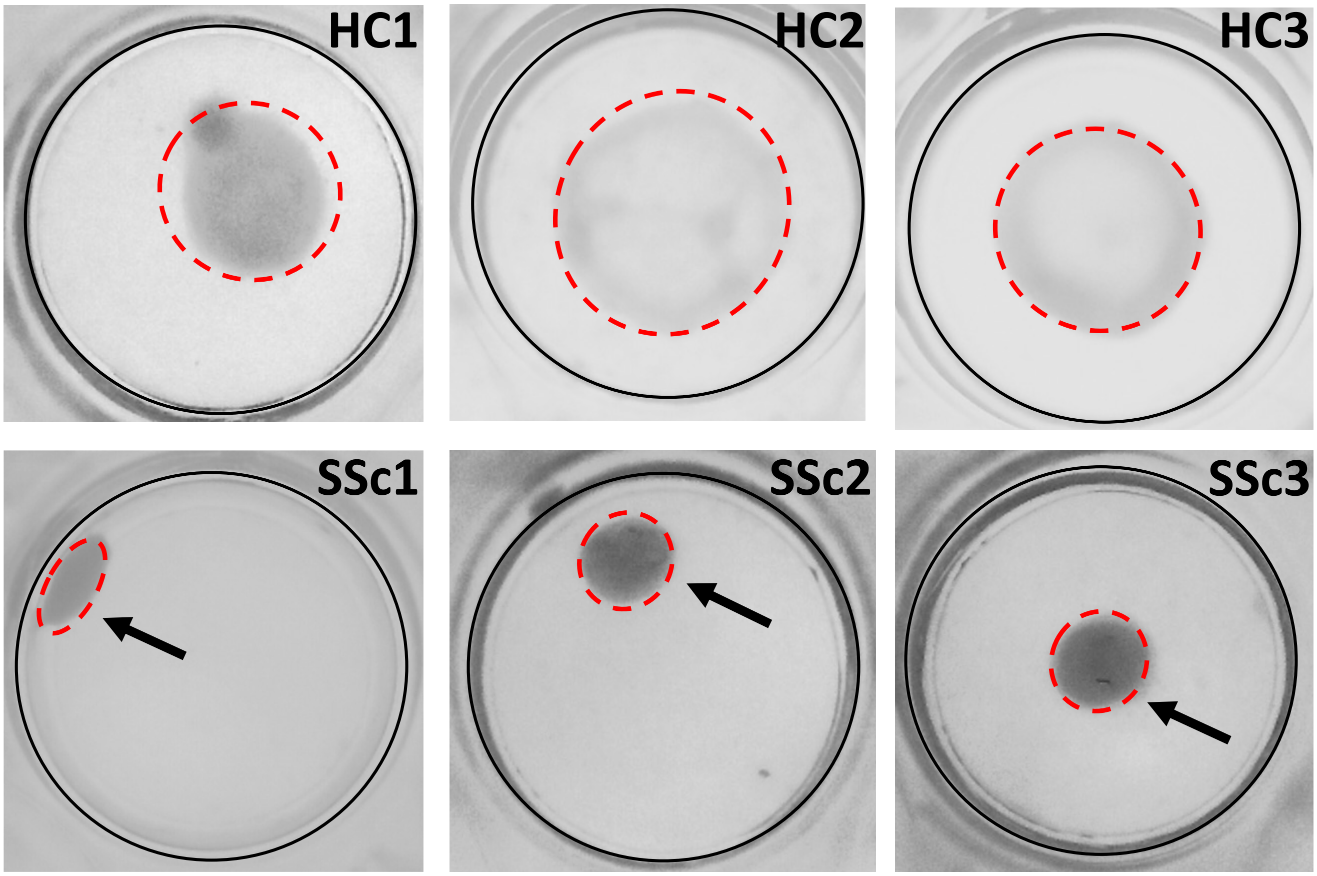


**Figure S1. Characterization of self-assembled skin equivalent tissues.** Self-assembly skin equivalent tissues were grown from autologous materials from three healthy control donors (HC1, HC2, HC3) and three systemic sclerosis patients (SSc1, SSc2, SSc3). Representative overhead images of tissues shown. Black line delimits the edges of the trans well insert. Red dotted line shows perimeter of tissue. Arrows point to contracted tissue in wells. HC = healthy control, SSc = systemic sclerosis

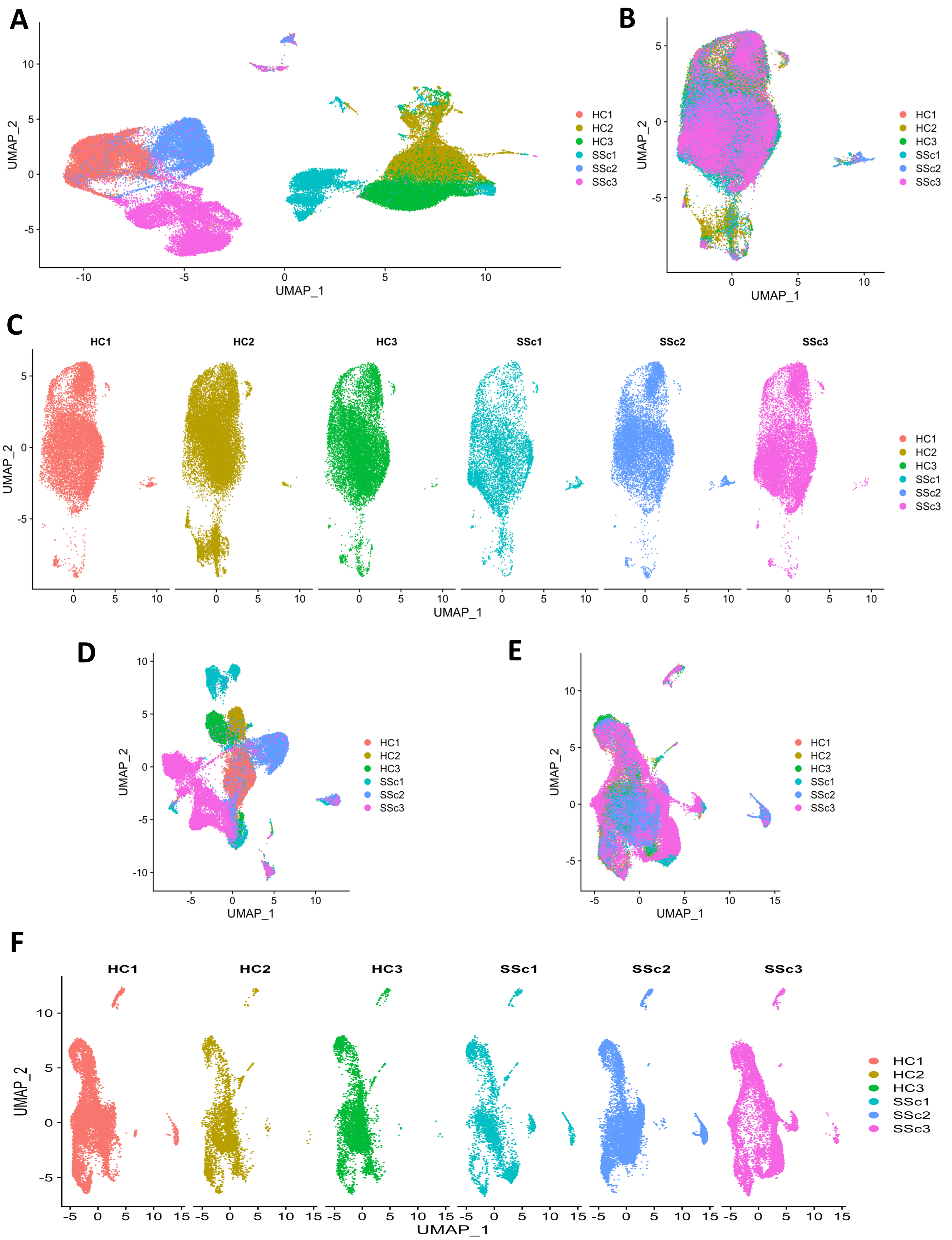


**Figure S2. Merged and integrated multiomic data at single-cell resolution.** A) Merged scRNA-seq data colored by sample. B) Integrated scRNA-seq data colored by sample. C) Integrated scRNA-seq data split by sample. D) Merged scATAC-seq data colored by sample. E) Integrated scATAC-seq data colored by sample. F) Integrated scATAC-seq data split by sample. HC = healthy control, SSc = systemic sclerosis

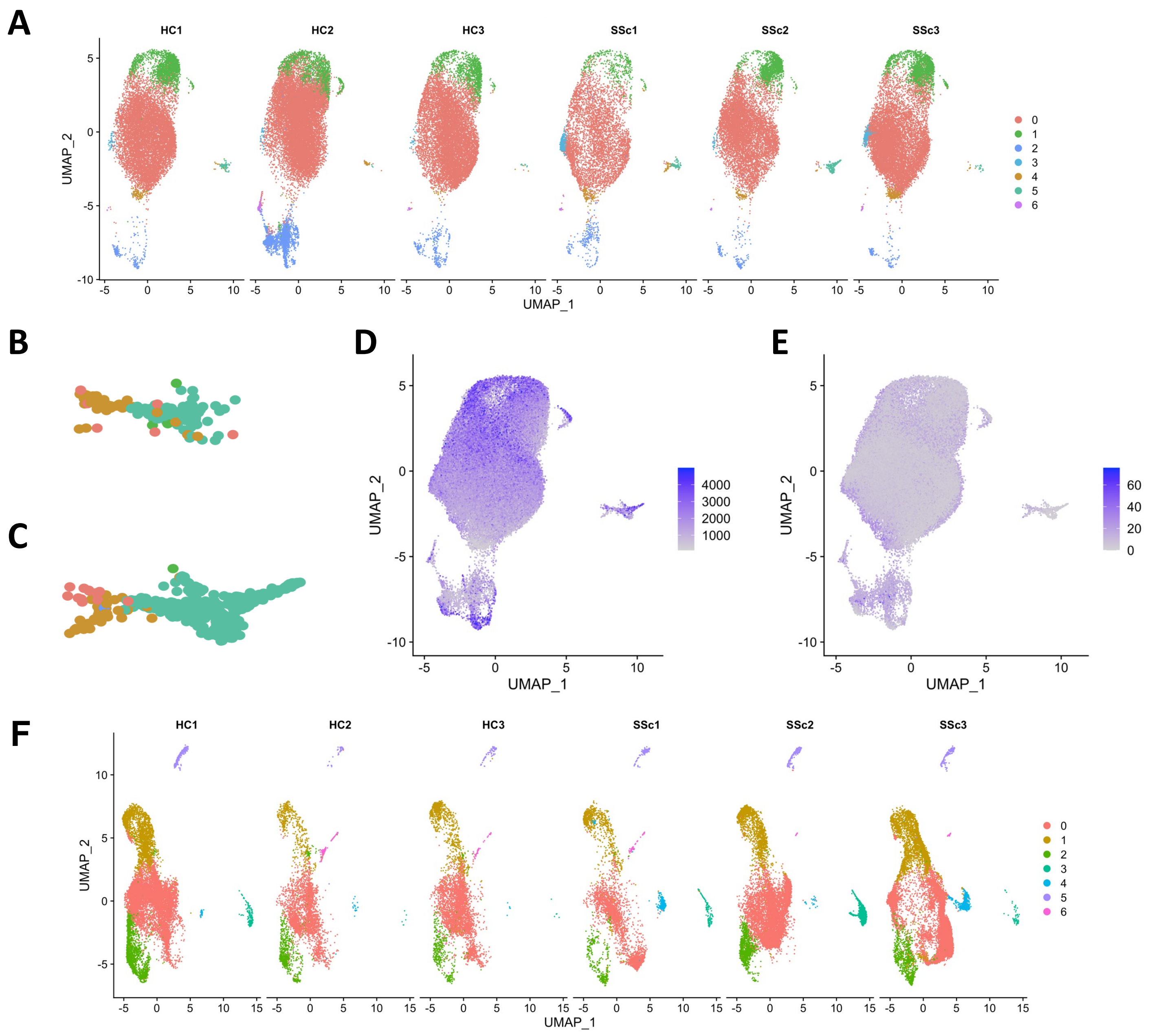


**Figure S3. Cluster determination and quality control for integrated data from single-cell multiomics of self-assembled skin equivalents.** A) UMAP projections of clustered transcriptomic data split by sample B-C) Close-up view of adjacent macrophage and fibrocyte-like cell clusters for both B) healthy control (HC) and C) systemic sclerosis (SSc) samples. D) Number of unique genes expressed for each cell plotted as a feature plot with increasing number of genes shown by increasing intensity of color. E) Percent of ribosomal genes plotted for each cell on a feature plot and increasing percentage denoted as more intense color. F) UMAP projections of clustered single-cell epigenomic data split by sample. HC = healthy control, SSc = systemic sclerosis

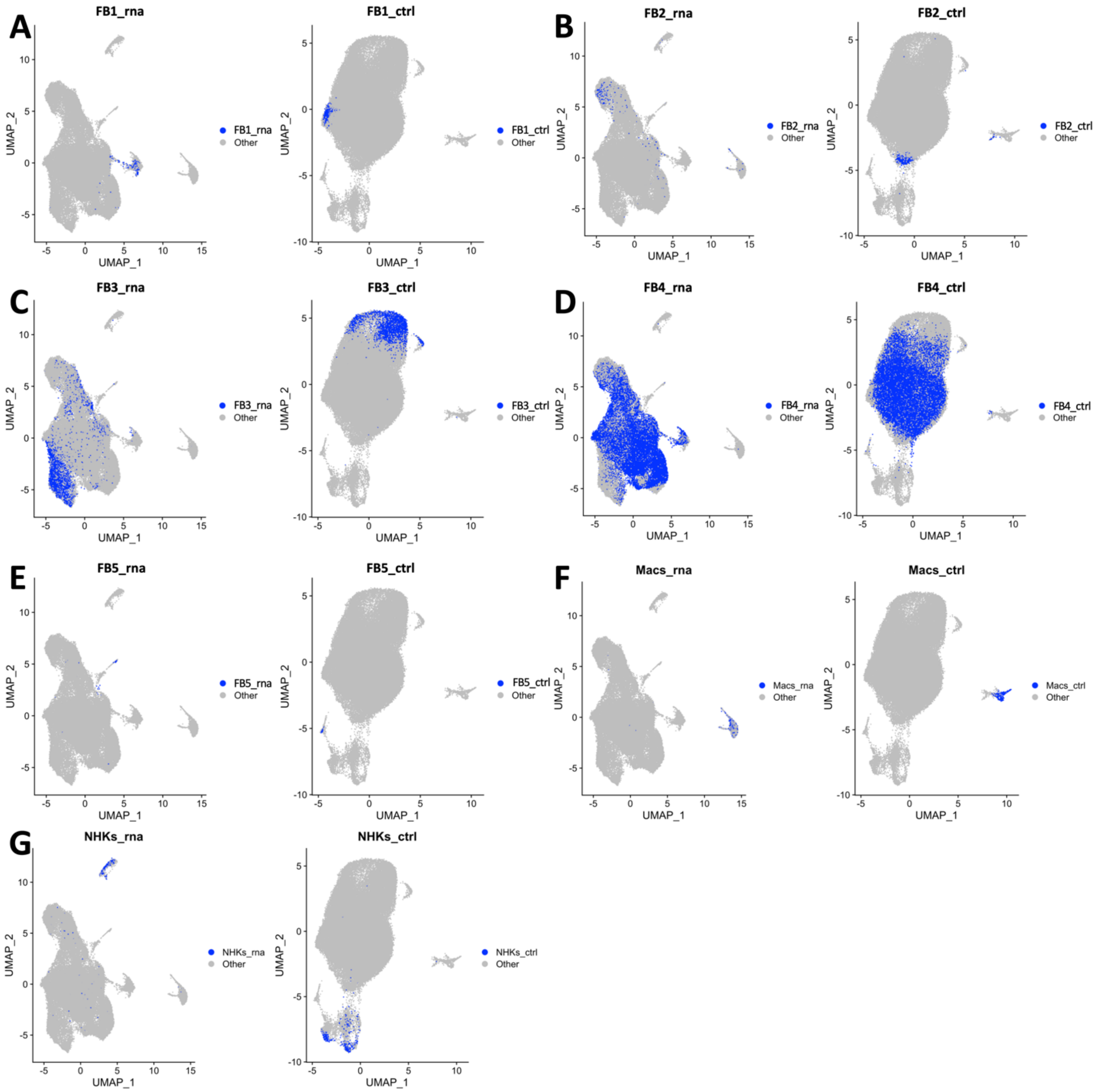


**Figure S4. Multiomic comparison of clusters from single-cell transcriptomic data compared to single-cell epigenomic data.** A-G) Intersecting multiome cells from transcriptomic data (right panel) were projected onto the corresponding cells in integrated epigenomic data (left panel). Macs = macrophages, NHKs = normal human keratinocytes

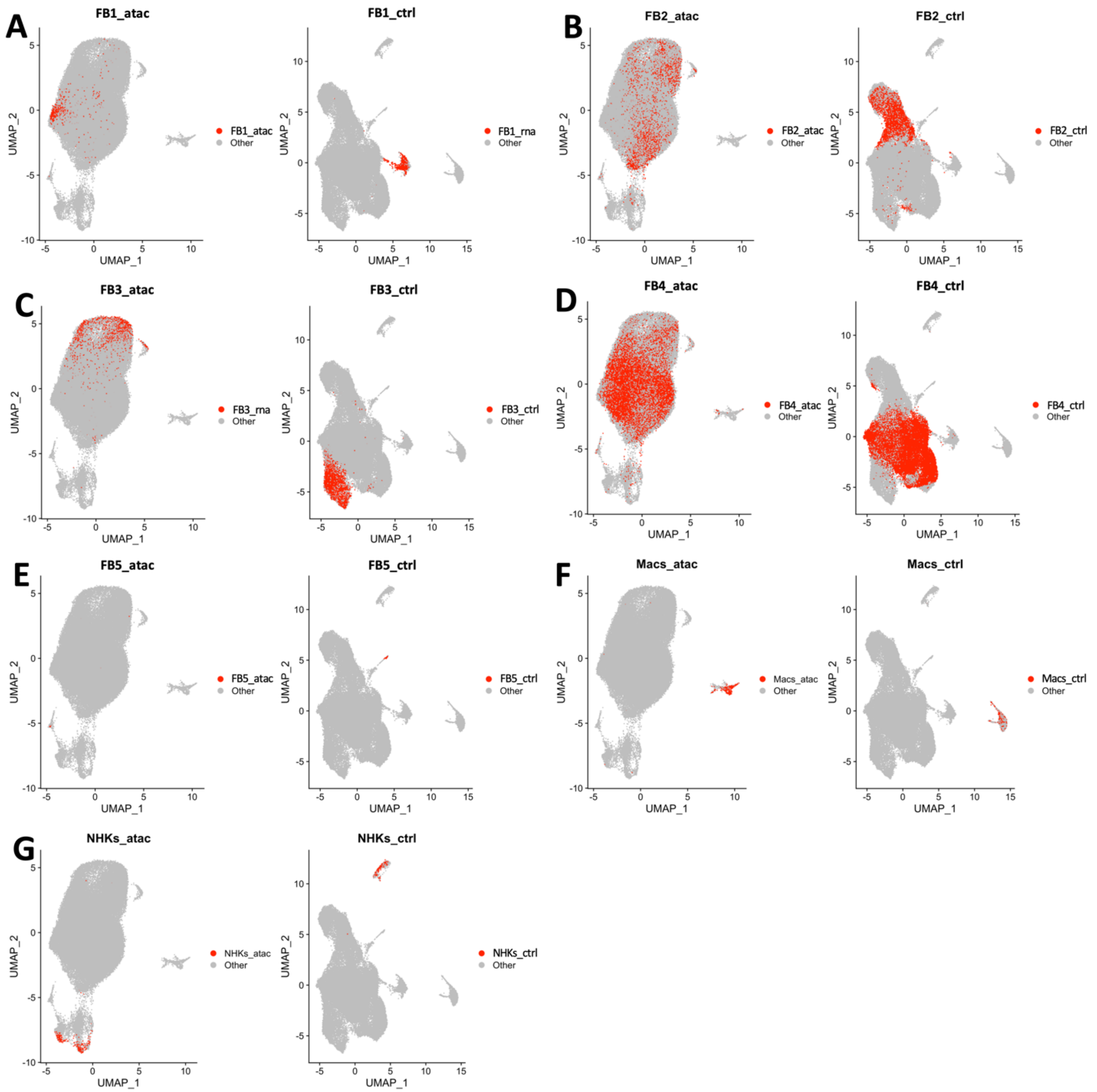


**Figure S5. Multiomic comparison of clusters from single-cell epigenomic data compared to single-cell transcriptomic data.** A-G) Intersecting multiome cells from epigenomic data (right panel) were projected onto the corresponding cells in integrated transcriptomic data (left panel). Macs = macrophages, NHKs = normal human keratinocytes

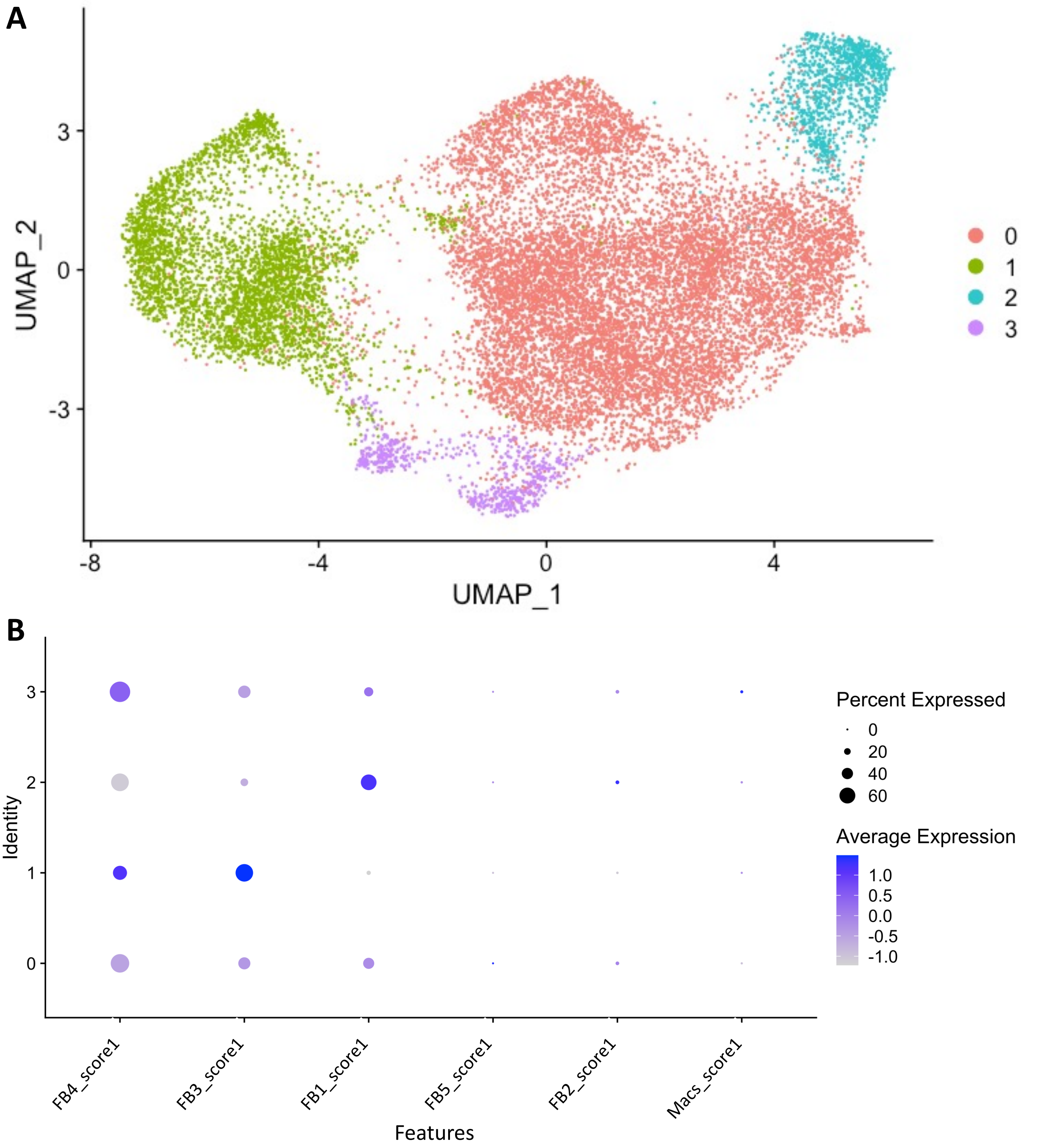


**Figure S6. Analysis of self-assembled stromal tissues in the absence of monocytes.** A) UMAP projection of clustered single-cell RNA-seq data from self-assembled stromal tissues grown in the absence of monocytes. One healthy control replicate and one SSc condition replicate. B) Fibroblast cluster score from full self-assembled stroma tissues containing monocytes applied to clusters from tissues with no monocytes added.

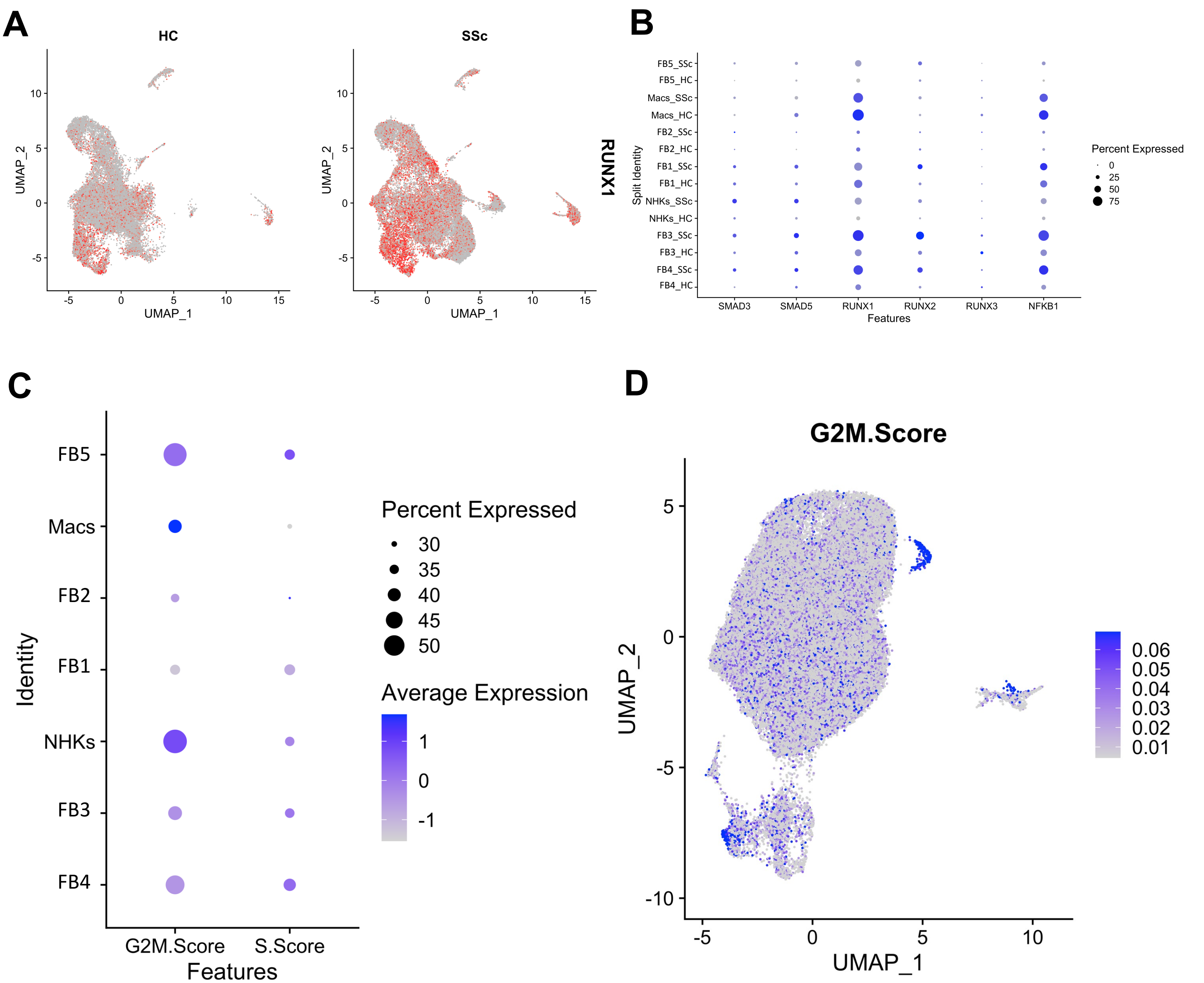


**Figure S7. Gene expression for select transcription factors (SMAD3/5, RUNX1/2/3, NFKB1) enriched in FB3 fibroblast population and cell cycle of 3D tissue clusters.** A) Feature plot for RUNX1 motif accessibility in scATAC-seq data split by disease state. B) Dot plot of single-cell gene expression for transcription factors determined to have enriched motifs in FB3 fibroblast population. Dot plot grouped by cluster and split by disease state. C) Dot plot showing combined G2 and M phase scores (G2M.Score) as well as S phase scores (S.Score) for each cluster. D) Feature plot showing strength of G2/M phase gene expression in cells across integrated single-cell transcriptomic data. HC = healthy control, SSc = systemic sclerosis, FB = fibroblast, Macs = macrophages, NHKs = normal human keratinocytes

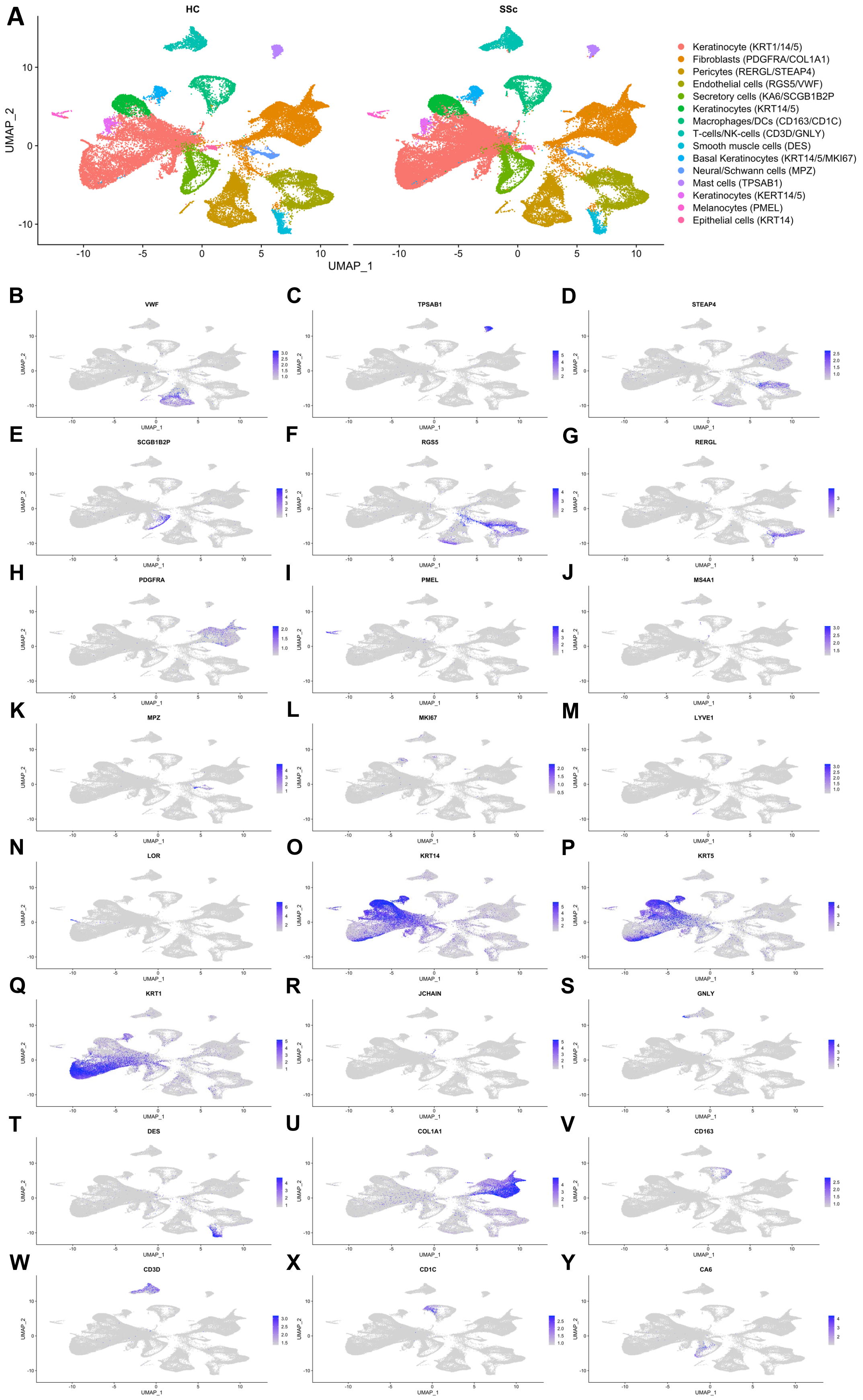


**Figure S8. Cell type assignment of clusters from integrated Tabib *et al.* single-cell RNA-seq data.** A) UMAP projection of integrated and clustered data with cell type labels based on gene expression of cell-type specific genes visualized in feature plots. B-Y) Feature plots of select cell-type specific genes in the integrated scRNA-seq data from human skin.

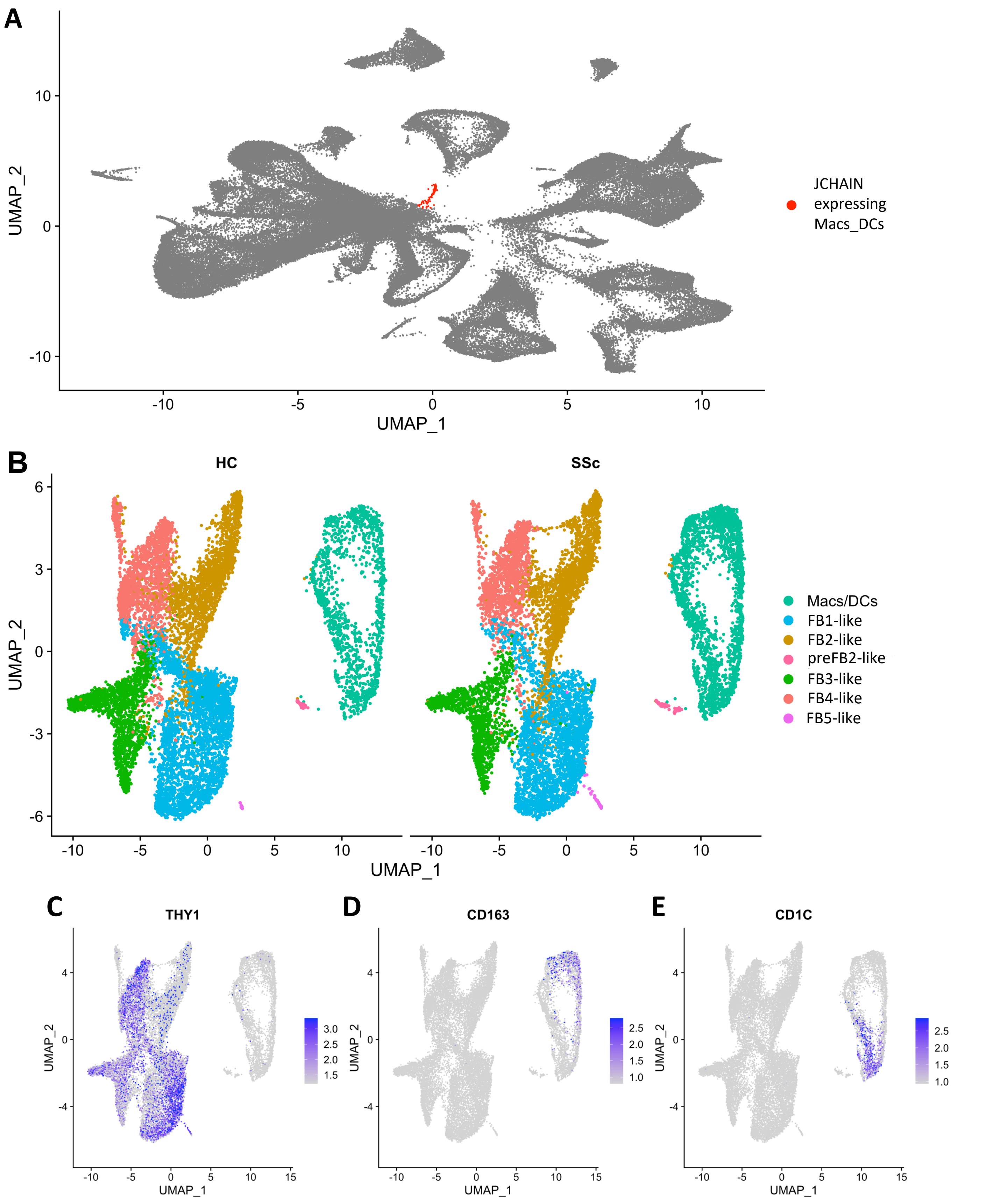


**Figure S9. Re-clustered fibroblast, macrophage, and dendritic cells from Tabib *et al.* single-cell RNA-seq.** A) UMAP projection of full integrated data showing JCHAIN expressing population of myeloid cells selected to include in re-clustered data. B) Integrated and clustered data. C) Feature plot of cell type markers for fibroblasts (THY1), macrophages (CD163), and dendritic cells (CD1C) projected on re-clustered data. HC = healthy control, SSc = systemic sclerosis

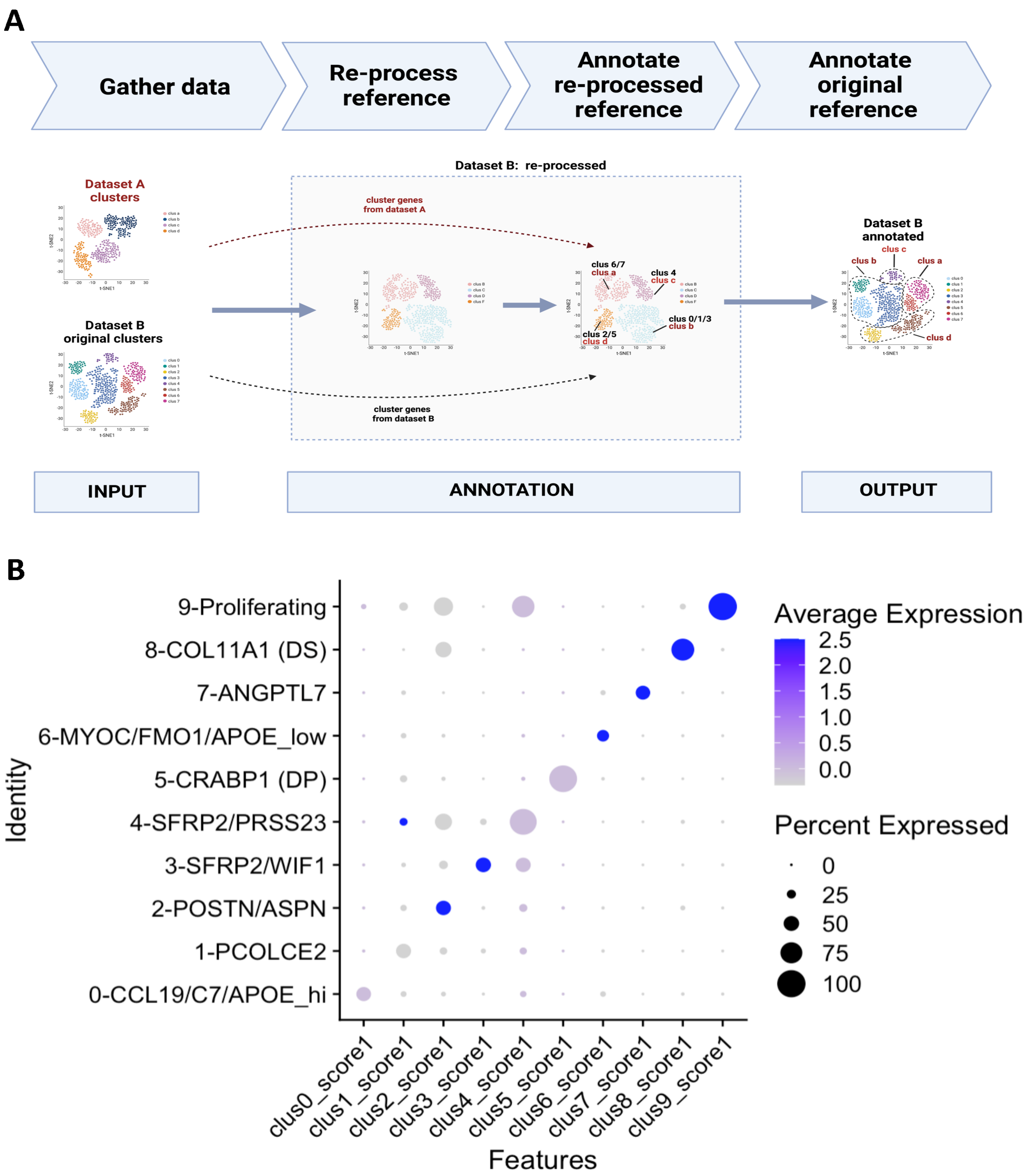


**Figure S10. Computational strategy for annotation of 3D tissue fibroblast clusters in skin data.** A) Diagrammatic representation of computational strategy used to identify analogous clusters between two different single-cell datasets. P) Evaluation of cluster scoring methodology. Dot plot using top 50 genes from each original fibroblast cluster applied to the original clustered fibroblast object from Tabib *et al.* publication.

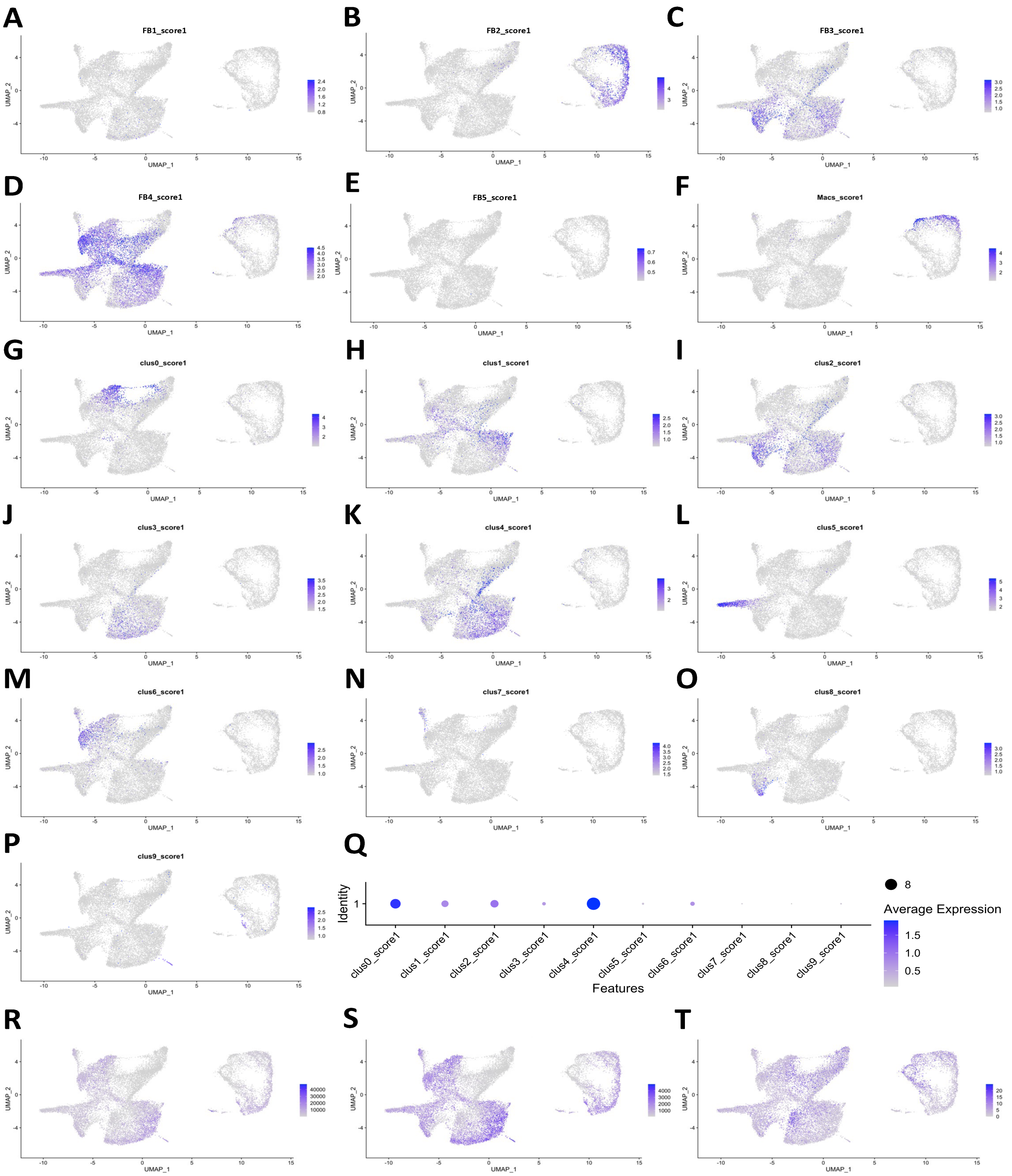


**Figure S11. Annotation of skin fibroblast clusters in re-processed and clustered scRNA-seq fibroblast data.** A-F) Feature plots of integrated Tabib *et al.* data with projected cluster scores obtained from top differentially expressed genes in single-cell transcriptomic clusters from self-assembled skin equivalent tissues. G-P) Feature plots of re-processed Tabib *et al.* data with projected cluster scores obtained from top 50 differentially expressed genes in original fibroblast clusters from publication. Q) Simple dot plot for cluster 1 of re-processed skin data which showed little expression for any of the original Tabib *et al.* fibroblast clusters. R-T) Feature plots of quality control metrics including number of total transcripts (R), number of unique genes (S), and number of mitochondrial transcripts per cell (T). HC = healthy control, SSc = systemic sclerosis, FB = fibroblast, Macs = macrophages, NHKs = normal human keratinocytes

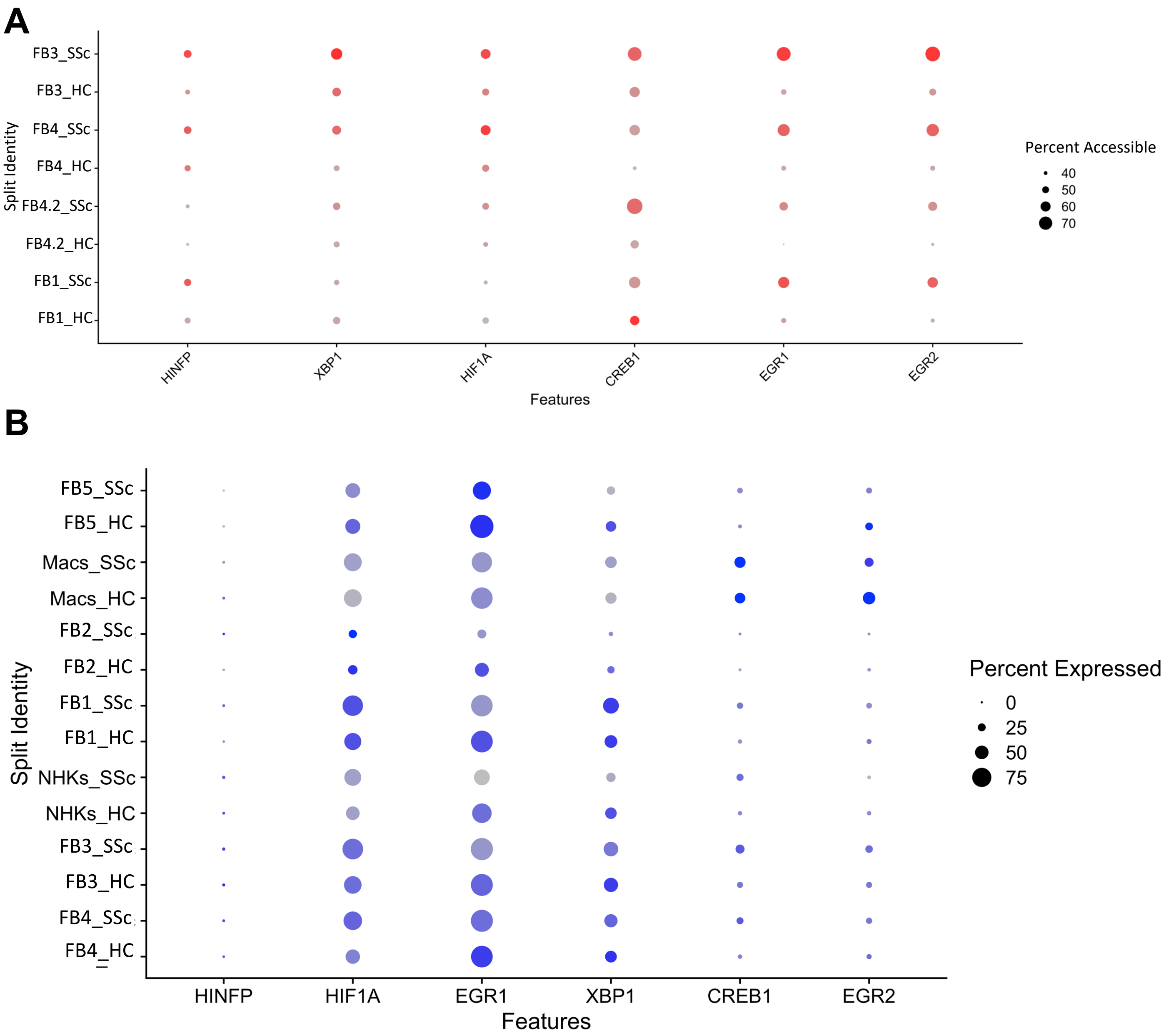


**Figure S12. Transcription factor motif enrichment in systemic sclerosis fibroblasts from single-cell ATAC-seq data from self-assembled skin equivalent tissues.** A) Dot plot of select SSc-enriched transcription factor motifs in fibroblast clusters split by disease state. B) Dot plot of gene expression for SSc-enriched factors in single-cell transcriptomic data grouped by cluster and split by disease state. HC = healthy control, SSc = systemic sclerosis, FB = fibroblast, Macs = macrophages, NHKs = normal human keratinocytes

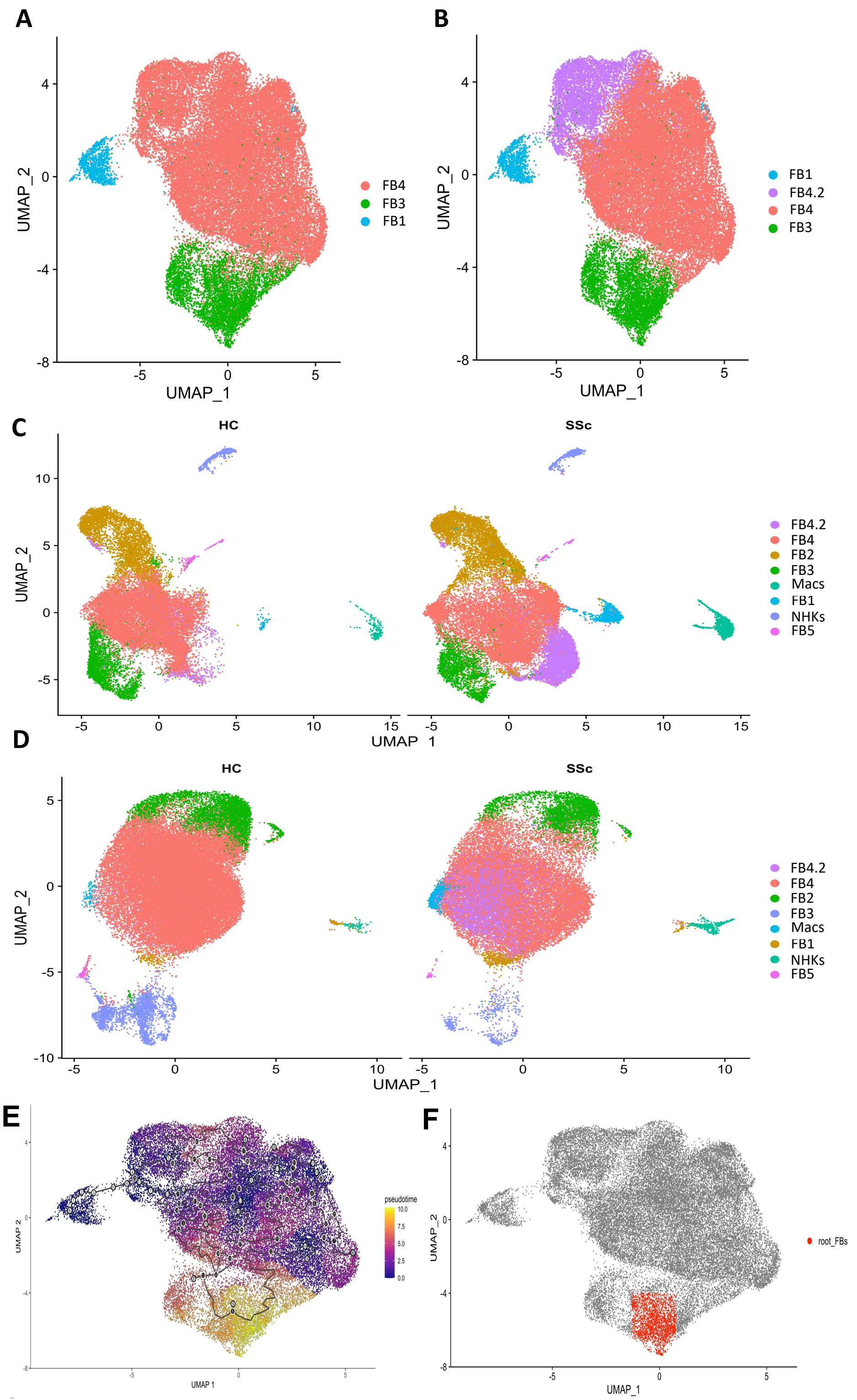


**Figure S13. Characterization of FB4.2 cluster of fibroblasts and pseudotime trajectory.** A) UMAP projection of re-clustered fibroblast clusters labeled and colored by original cluster assignment from full integrated object. B) UMAP projection of clusters colored and labeled according to new clusters determined from re-clustering including cluster FB4.2 (purple). C) All scATAC-seq cells from self-assembled skin equivalent (saSE) tissues with cells from FB4.2 cluster highlighted (purple). D) All scRNA-seq data from saSE tissues with cells from FB4.2 cluster highlighted (purple). E) UMAP projection of pseudotime trajectory of fibroblasts when using the FB1 population as the starting population. F) UMAP projection showing selected root cells (red) for pseudotime trajectory predicting path of FB1 differentiation. HC = healthy control, SSc = systemic sclerosis, FB = fibroblast, Macs = macrophages, NHKs = normal human keratinocytes

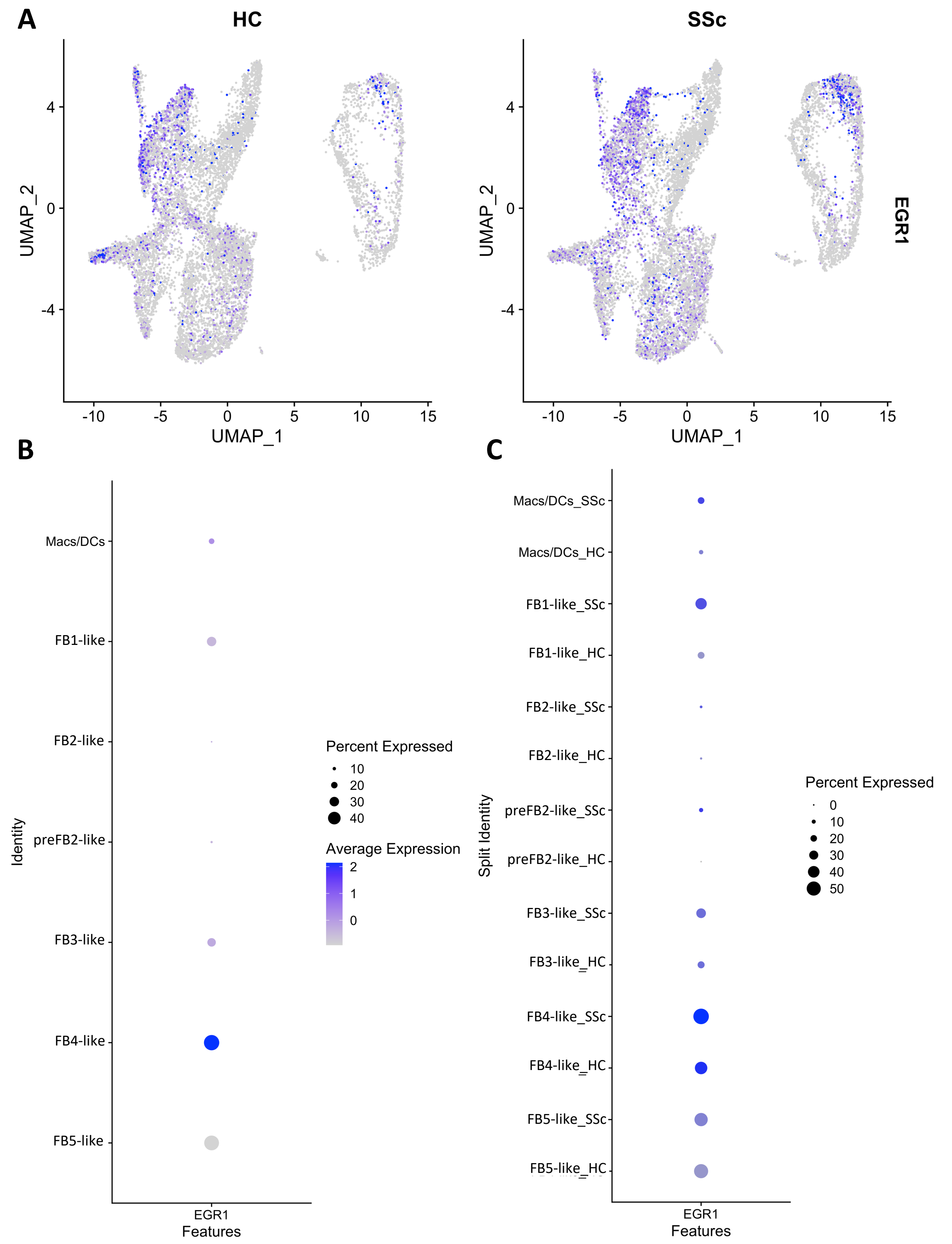


**Figure S14. EGR1 Expression in single-cell RNA-seq data from human skin (Tabib *et al.*).** A) Feature plot of fibroblasts, macrophages, and dendritic cells with EGR1 expression in blue. B) Dot plot of EGR1 gene expression in fibroblast clusters from Tabib *et al.* C) Dot plot split by disease state. HC = healthy control, SSc = systemic sclerosis, FB = fibroblast, Macs = macrophages, DCs = dendritic cells, NHKs = normal human keratinocytes

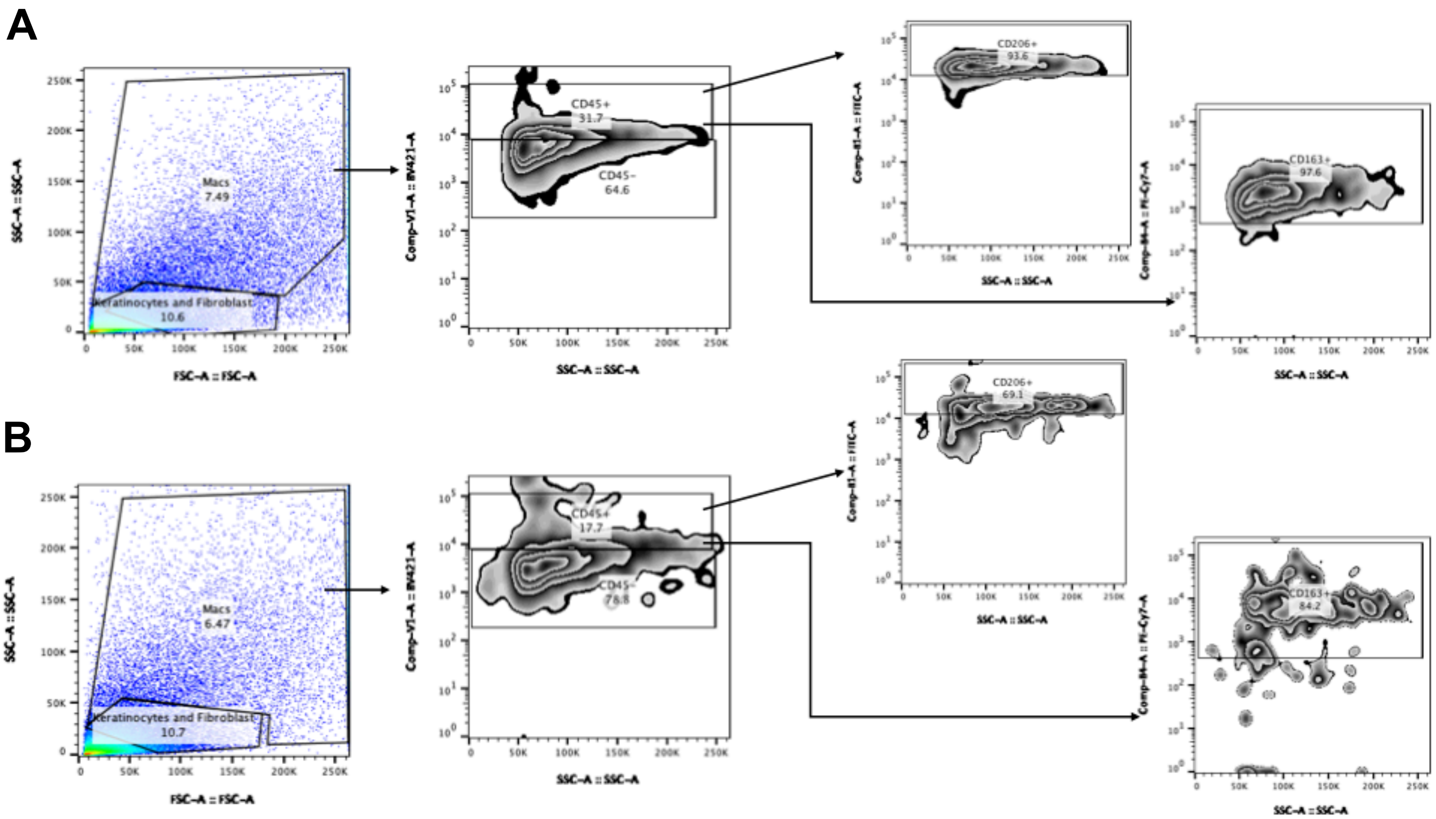


**Figure S15. Flow analysis of cells from representative self-assembled stromal tissues.** Flow analysis of dissociated cells from representative self-assembled skin-equivalent tissues cultured using A) systemic sclerosis and B) healthy control cells. Cell size and granularity were used to isolate macrophages. CD45 + cells were selected and resulting population was analyzed for percent CD206+ and CD163+ cells which is a marker of IL-4 activation.

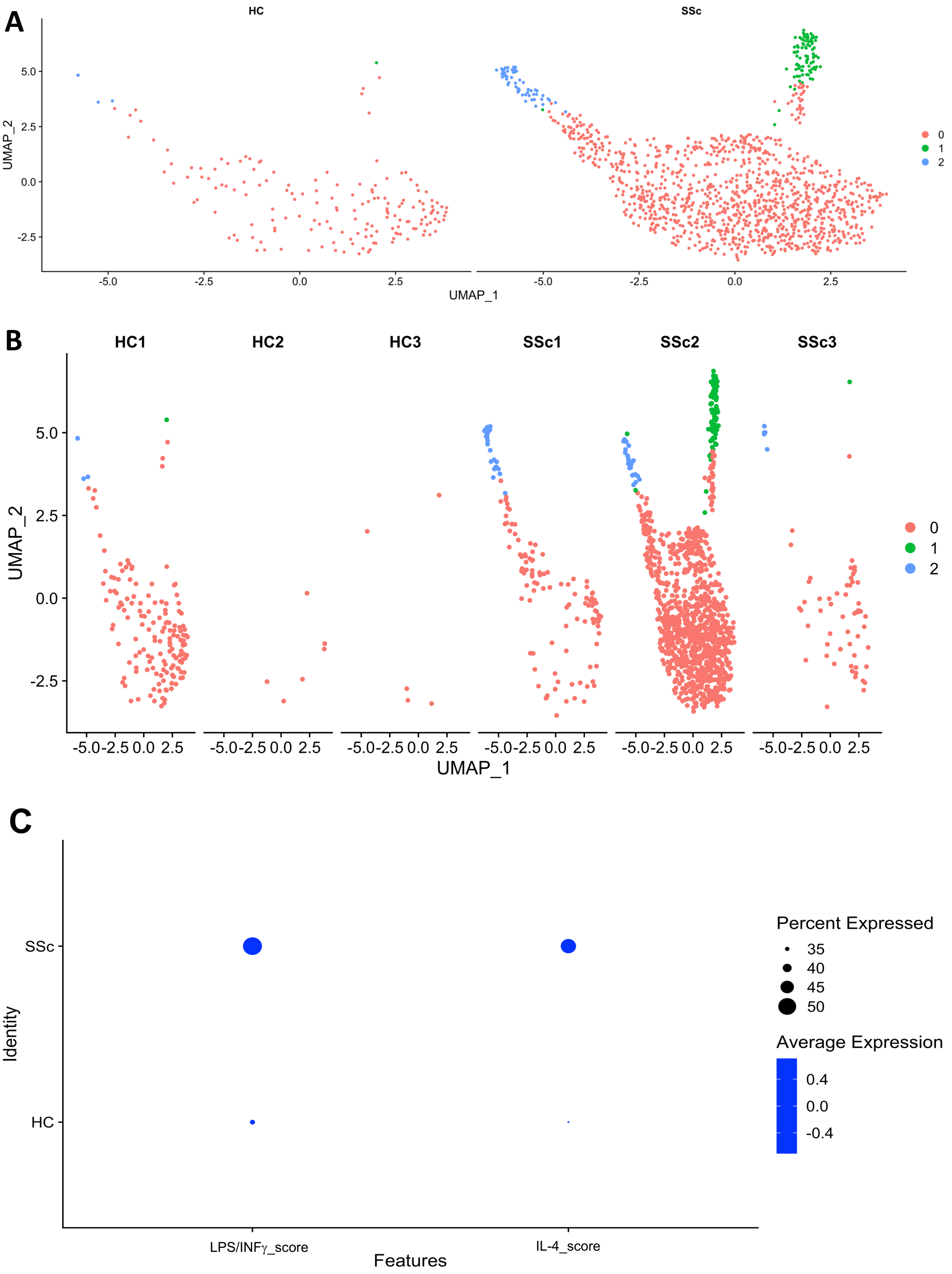


**Figure S16. Single-cell ATAC-seq Macrophage subsets in self-assembled skin equivalent tissues.** A) UMAP projections of clustered macrophages split by disease state. B) UMAP projections of macrophages split by sample. C) Dot plot of LPS/INFγ and IL-4 activation scores in macrophages split by disease state.

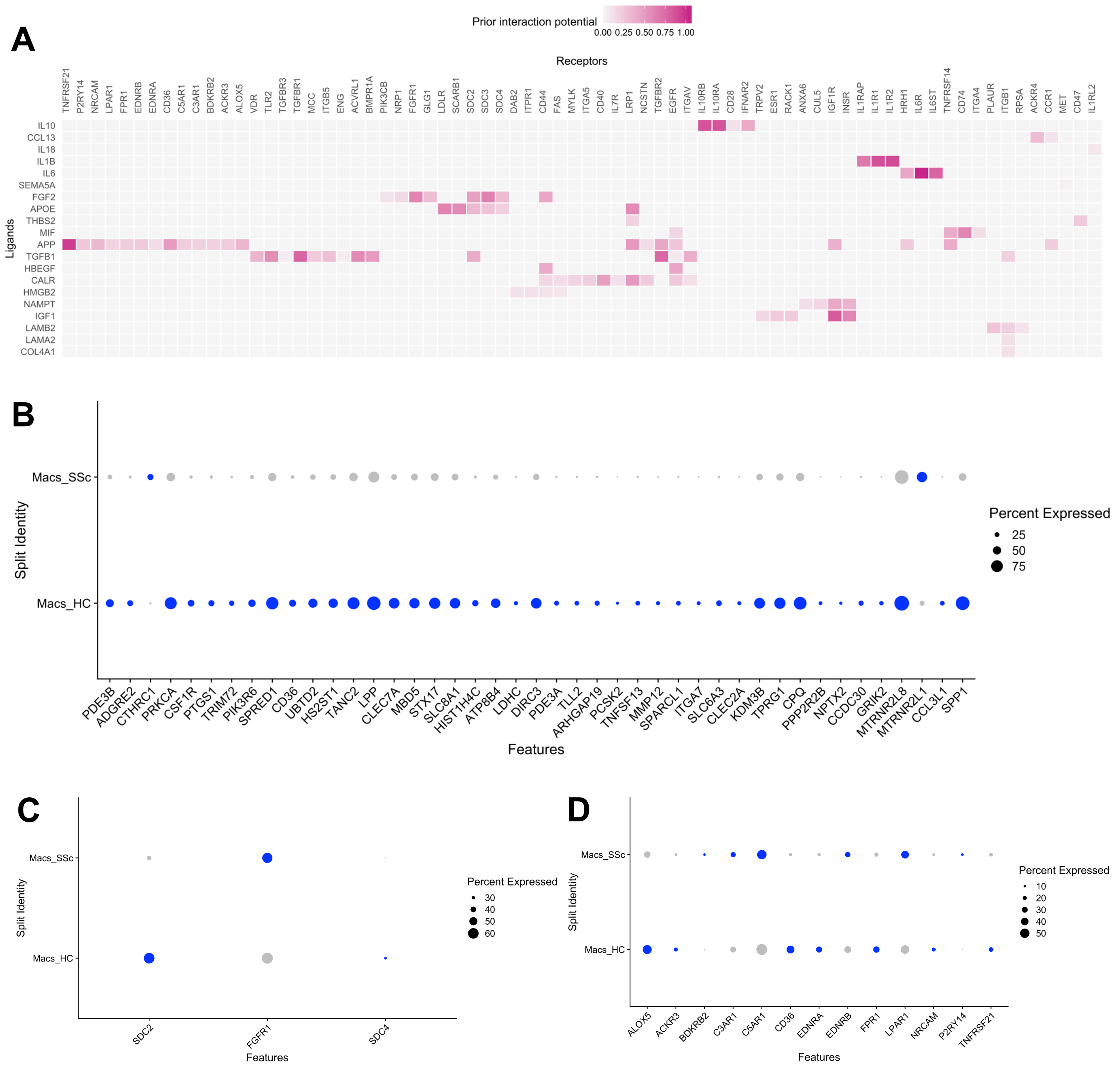


**Figure S17. Ligand-receptor interaction potential of ligands from fibroblasts, macrophage target gene expression, and macrophage receptor expression from self-assembled skin equivalent tissues.** A) Predicted ligand-receptor interactions for top fibroblast ligands determined from NicheNet analysis. B) Dot plot of macrophage target genes as determined by NicheNet analysis split by disease state. C) Dot plot of FGF2 receptors in macrophages split by disease state. D) Dot plot of APP receptors in macrophages split by disease state. HC = healthy control, SSc = systemic sclerosis, Macs = macrophages
