## Supplemental Methods for "Single-cell epigenomic dysregulation of Systemic Sclerosis fibroblasts via CREB1/EGR1 axis in self-assembled human skin equivalents"

***Pre-processing of scRNA-seq Data***

Data were normalized, integrated, and scaled using a regression to remove batch effects. Clusters determined via ClustTree were relatively evenly distributed across samples from the same disease state (Fig. S5A).

Poor quality cells were filtered from each sample based on number of features (> 100, <5000), minimum number of cells (number = 3), and percent mitochondrial reads (less than 25%). We used more lenient mitochondrial filtering to avoid removing macrophages which are highly metabolic and as such had high mitochondrial gene expression. Samples were independently log normalized and variable features were identified prior to integration (dimensions = 1:50). Although we initially attempted to normalize and integrate transcriptional data using single-cell transform, it appeared to overcorrect the data contributing to difficulty in identifying clusters (data not shown). We chose log normalization since it didn't overcorrect and retained the greatest cluster diversity allowing for more sensitive detection of clusters which matched those identified in the scATAC-seq data. After merging and integrating the data was scaled and corrected for batch using a regression model. Integration resulted in an even distribution of cell transcriptomic profiles across the six samples (Fig. S2B-C). An elbow plot of principal components was used to determine appropriate dimensions (dimensions = 1:30). Cluster resolution and cluster enrichment were determined using clustree (resolution = 0.09). Cell proportion differences were calculated as both log fold change (gtools v3.9.3) and odds ratio (questionr v0.7.7) and visualized in ggplot2. Top differentially expressed genes in each cluster were determined using FindMarkers tool in Seurat, comparing each cluster to all other clusters and filtered (p <0.05). Markers were ranked based on fold enrichment and used as input on gprofiler to determine GO terms associated with each cluster.

The CellCycleScoring command in Seurat was used to add cell cycle information by scoring each cell based on the gene expression of G1, G2, S, and M phase genes.

### *Pre-processing of scATAC-seq Data*

For chromatin data, low quality cells were filtered based on outliers in nucleosome enrichment, transcription start site enrichment, and other key quality metrics. The first dimension was excluded from all downstream analysis due to high correlation with sequencing depth and in compliance with standard recommended protocol for analyzing scATAC-seq data. Samples were normalized, reduced, merged (dimensions = 2:50), and integrated (dimensions = 2:50) using standard established methods. Integration of scATAC-seq samples resulted in even distribution of cells across samples (Fig. S5E-F) Results were visualized on UMAP projections using dimensions determined by the elbow plot of reduction data (dimensions = 2:30). Cluster resolution was determined using the clustree package (resolution = 0.08). Cell proportion differences were calculated as both log fold change (gtools v3.9.3) and odds ratio (questionr v0.7.7) and visualized in ggplot2. Transcription factor enrichment for each cluster as compared to all other clusters was determined using FindMarkers and FindMotifs function in Signac. Peaks were filtered by fold-change prior to determining motifs (average log2 fold change > 0.2, but > 0.1 for FB4 and FB2 which had low fold change). Resulting motifs were filtered ($p < 0.05$) and ranked by fold enrichment. FeaturePlot and DotPlot functions in Seurat were used to visualize markers of interest.

### *RNA/ATAC Multiome Comparison*

Using Seurat, we extracted cell names separately for multiome cells from each cluster in both integrated modalities (RNA and ATAC). We found the intersection of cell names for each cluster from both ATAC and RNA modalities. Cell names were assigned to a new ident and plotted to their respective counterpart (i.e., intersecting multiome cells detected in the ATAC cluster were plotted on the RNA integrated dataset and vice versa). Cells names were also plotted back to the original modality as a control.

***RNA/ATAC Integration***

Predicted gene expression was calculated in ATAC integrated object using Ensemble annotation and Seurat GeneActivity command. Predicted expression was normalized and scaled. FindTransferAnchors was used to find anchor cells with similar gene expression in RNA integrated object with ATAC object as the reference object. Cell labels were transferred to the RNA object based on predicted anchors and results were plotted using UMAP visualization.

***Analysis of scRNA-seq Data from Human Skin***

Single cell RNA-seq data from the publication by Tabib et al. (5) was downloaded from Gene Expression Omnibus (GSE138669). Data were pre-processed as described previously in section "Pre-processing of scRNA-seq Data"**.** The high compute cluster Discovery at Dartmouth College was used for integration of data. It should be noted that data was of low read depth (majority of cells <50 features per cell), so a large number of cells were lost in filtering. More lenient filtering retained more cells but resulted in difficulty visualizing dimensionally reduced data (i.e., feature space was too low for effective dimensional reduction in UMAP plots). Clusters were determined within the following parameters: dimensions = 1:50, resolution = 0.07. Cell type was assigned to each cluster using gene expression distribution of cell type specific markers shown in original publication. The fibroblasts, macrophage/DC, and a small population of JCHAINexpressing macrophages/DCs were isolated for additional clustering (dimensions: 1:50, resolution: 0.09). Fold change and odds ratio were determined as previously described in section "Pre-processing of scRNA-seq Data".

### *Self-Assembled Stromal Tissues without Monocytes*

Data were analyzed as described previously in section "Pre-processing of scRNA-seq Data" (dimensions = 1:30, resolution = 0.1). Top ten DE genes from each cluster in the saSE tissue data were used to create fibroblast subset scores using the Seurat AddModuleScore command and plotted using FeaturePlots.

### *Annotation of Fibroblast Subclusters Using Human Skin Reference*

We followed the four-step strategy outlined in Supplemental Figure S10A. First, we gathered our reference (human skin) and novel (saSE tissue) datasets. For the reference dataset we gathered the raw files from the Gene Expression Omnibus (GSE138669) and top differentially expressed genes for each cluster from the supplemental tables for the Tabib et al publication (5). For our novel dataset, we gathered information on processing parameters (filter cutoffs, dimensions, etc.) and top differentially expressed genes for each cluster. Secondly, we re-processed the reference dataset from the Tabib et al. publication, isolated fibroblasts and macrophages, and re-clustered as described for our saSE tissues. By re-processing and clustering the skin data using similar parameters as applied to the saSE tissue data, we controlled for potential confounders due to differences in analysis pipeline, creating a common reference for both datasets. Thirdly, we annotated this common reference using the top differentially expressed genes from both the original Tabib et al. fibroblast clusters and our saSE tissue fibroblast clusters. The top fifty DE genes from each cluster in the saSE tissue data were used to create fibroblast subset scores using the Seurat AddModuleScore command and plotted using FeaturePlots and DotPlots. Clusters containing the highest module score as determined by the dot plots were annotated accordingly. Some module scores with genes containing low or sparse expression were plotted separately from those with higher expression to differentiate between clusters more effectively. Fourthly, we manually transferred labels from the common reference to the coordinating clusters in our novel saSE tissue object. Thus, we were able to identify analogous clusters across the two datasets using publicly available data for our reference dataset.

***Fibroblast Cell Re-clustering and Pseudotime Analysis***

FB1, FB3, and FB4 cells in ATAC integrated object were isolated and re-clustered (dimensions = 2:20, resolution = 0.1). When FB2 fibroblasts were included, no additional information was gained in pseudotime analysis (data not shown). Pseudotime was performed using standard monocle3 analysis pipeline for Seurat samples. First pseudotime was conducted backwards using FB1 population as the starting population. Based on this backwards trajectory, FB3 cells were selected using the SelectCells function in Seurat for setting root of forward trajectory. Similar analysis was performed in RNA data using the same clusters and multiome cells from root cells selected in ATAC object to guide selection of root for trajectory.

***Macrophage Re-clustering and Analysis***

Macrophages were isolated and re-clustered as described for pre-processing of scATAC-seq data (dimensions = 2:30, resolution = 0.06). Cell scores were assigned to each cell using pre-designated LPS/INFγ or IL-4 associated transcription factor motifs (95-98). LPS/INFγ score included IRF3, IRF5, IRF7, IRF9, STAT1, and P65 TF motifs. IL-4 score included IRF4, STAT3, PPARG, KLF4, HIF1A, and P50/NFKB1 TF motifs. Fold enrichment and odds ratio were calculated as described for processing of scATAC-seq and scRNA-seq data. Differentially accessible motifs between all SSc Macrophages and all HC macrophages were determined as described previously for scATAC-seq data in section "Pre-processing of scATAC-seq Data".

***Ligand-Receptor Analysis***

Sender cells were defined as all fibroblasts (FB1, FB2, FB3, FB4, and FB5) and receiver cells were defined as the macrophage cluster. All preset parameters were used except that the "RNA" assay was specified as the data slot and number of L-R pairs to use from the ligand target matrix was increased to 3000 to ensure that all significant upregulated target genes were included in final analysis.
